# Nitrous oxide respiration in acidophilic methanotrophs

**DOI:** 10.1101/2024.01.15.574570

**Authors:** Samuel Imisi Awala, Joo-Han Gwak, Yongman Kim, Man-Young Jung, Peter. F. Dunfield, Michael Wagner, Sung-Keun Rhee

**Author notes:** Sung-Keun Rhee **Email:**. S.I.A and J-H.G contributed equally to this work.

## Abstract

Methanotrophic bacteria mitigate methane (CH_4_) emissions from natural environments. Although aerobic methanotrophs are considered strict aerobes, they are often highly abundant in extremely hypoxic and even anoxic environments. Despite the presence of denitrification genes, it remains to be verified whether denitrification contributes to their growth. Here, we revealed that two acidophilic methanotrophs encoding N_2_O reductase (clade I and type II nosZ, respectively): *Methylocella tundrae* T4 and *Methylacidiphilum caldifontis* IT6, respired N_2_O and grew anaerobically on diverse non-methane substrates, including methanol, C-C substrates, and hydrogen. However, NO_3_ ^−^ and NO_2_ ^−^ could be reduced during methanol oxidation in *Methylocella tundrae* T4 and *Methylocella silvestris* BL2 without significantly increasing cell biomass. The lack of growth on methanol + NO_3_^−^ or NO_2_^−^ was likely due to the production of toxic reactive nitrogen species and C1 metabolites. However, the oxidation of pyruvate, a C3 electron donor, combined with NO_3_^−^ or NO_2_^−^ reduction resulted in anaerobic growth of *Methylocella tundrae* T4 and *Methylocella silvestris* BL2. In the extreme acidophile, *Methylacidiphilum caldifontis* IT6, N_2_O respiration supported cell growth at an extremely acidic pH of 2.0. In *Methylocella tundrae* T4, simultaneous consumption of N_2_O and CH_4_ was observed in suboxic conditions, both in microrespirometry and growth experiments, indicating the robustness of its N_2_O reductase activity in the presence of O_2_. Furthermore, CH_4_ oxidation per O_2_ reduced in O_2_-limiting conditions increased when N_2_O was added, indicating that cells of T4 can direct more O_2_ towards methane monooxygenase when respiring N_2_O as a terminal electron acceptor. Upregulation of *nosZ* and distinct repertories of methanol dehydrogenase-encoding genes (XoxF- and MxaFI-type) in *Methylocella tundrae* T4 cells grown anaerobically on methanol with N_2_O as the sole electron acceptor indicated adaptation mechanisms to anoxia. Our findings demonstrate that some methanotrophs can respire N_2_O independently or in tandem with O_2_, significantly expanding their potential ecological niche and paving the way for enhanced growth and survival in dynamic environments. This metabolic capability has application potential for simultaneously mitigating the emissions of the key greenhouse gases, CO_2_, CH_4,_ and N_2_O, from natural and engineered environments.

## Introduction

Anthropogenic emissions of greenhouse gases (GHGs)—primarily carbon dioxide (CO_2_), methane (CH_4_), and nitrous oxide (N_2_O)—are responsible for a historically rapid increase in Earth’s average annual temperature of more than 0.2°C per decade ^1, 2^. In addition to achieving net-zero CO_2_ emissions by 2050, significant reductions in the emissions of other GHGs including CH_4_ and N_2_O are now critically needed. Compared to CO_2_, the warming effect of CH_4_ is around 28 times greater ^3^. However, its much shorter mean lifetime of approximately 12–13 years ^4^ provides an additional opportunity to mitigate future climate change. Like CO_2_, N_2_O—the third most important GHG—has a long half-life (roughly 120 years) in the atmosphere ^5^, and its warming potential is about 300 times greater than CO_2_ over a 100-year time scale ^1^. In addition, N_2_O is a major cause of ozone depletion in the stratosphere ^6, 7^.

Although human activities are by far the most important reason for the unprecedented rise in atmospheric GHGs ^8^, microbial activities directly play a significant role in this rise ^9–12^. GHG net accumulation is regulated by the biogeochemical source-sink dynamics of GHGs exchanged between terrestrial, marine, and atmospheric reservoirs ^8^. GHG production and consumption in both natural and anthropogenic ecosystems are driven primarily by microbes ^9–12^. Methane fluxes in natural environments are controlled by activities of methane-producing (methanogenic) and methane-consuming (methanotrophic) microorganisms. It is estimated that 69% of the atmospheric CH_4_ budget originates from microbial activities (methanogenesis) while about 50-90% of the produced CH_4_ is oxidized by methanotrophs before reaching the atmosphere ^13–15^.

Microbes can oxidize methane under aerobic and anaerobic conditions. Aerobic methanotrophs oxidize methane to methanol by employing either particulate methane monooxygenases (pMMO) or soluble methane monooxygenases (sMMO) ^16–18^. There are two ways in which aerobic methanotrophs use oxygen ^19^: as the terminal electron acceptor of aerobic respiration and for methane activation via the methane monooxygenase ^16–18^. Under strictly anoxic conditions, anaerobic methanotrophic microorganisms mitigate CH_4_ emissions by oxidizing methane with alternative terminal electron acceptors including nitrate, Fe^3+^, Mn^4+^, sulfate, and humic acid using reverse methanogenesis pathways ^20–23^. Furthermore, intra-aerobic metabolism in the nitrite-dependent anaerobic methane-oxidizing bacterium ‘*Candidatus* Methylomirabilis oxyfera’ using pMMO was reported ^24^.

Interestingly, the genomes of some aerobic methanotrophs encode denitrification enzymes including nitrate (NO_3_^−^), nitrite (NO_2_^−^), nitric oxide (NO), and N_2_O reductases ^25–29^. Surprisingly, however, none of the methanotroph genomes or MAGs known to date encode a complete set of denitrification genes (Supplementary Table 1). Kits and colleagues ^26, 27^ demonstrated that aerobic methanotrophs can couple NO_3_^−^ and NO_2_^−^ reduction to the oxidation of methane and other electron donors, including methanol, formaldehyde, formate, ethane, ethanol, and ammonia in suboxic conditions. However, whether these aerobic methanotrophs are capable of anaerobic growth with NO_3_^−^ and NO_2_^−^ as terminal electron acceptors remain to be seen.

More than two-thirds of N_2_O emissions arise from bacterial and fungal denitrification and nitrification processes in soils ^9, 30, 31^. N_2_O emissions are a major concern in acidic environments due to the high production of N_2_O via abiotic reactions and the inhibition of biological N_2_O reduction ^32, 33^. Although multiple sources of N_2_O exist ^9, 30^, there is only one known sink for N_2_O in the biosphere—the microbial reduction of N_2_O to N_2_, catalyzed by a copper-dependent enzyme, N_2_O reductase (N_2_OR) encoded by *nosZ* ^34, 35^. The NosZ enzymes found in prokaryotes are phylogenetically classified into two clades: the canonical NosZ (clade I NosZ), found mostly in denitrifiers ^36^, and the recently described *c*NosZ (clade II NosZ) ^37^, which has an additional *c*-type heme domain at the C terminus, found commonly in non-denitrifiers ^37, 38^. Thus, bacteria and archaea harboring the *nosZ*-type genes—in particular those classified as incomplete- or non-denitrifiers—are receiving increasing attention in the search to combat N_2_O emissions ^38^. Previous studies have reported the presence of the *nosZ* gene in the aerobic methanotrophs, *Methylocystis* sp. SC2 ^39^ and *Methylocella tundrae* ^40, 41^: further genomic analysis from this study suggests that this enzyme is present in some other aerobic methanotrophs, too (Supplementary Table 1). Pure culture studies have unequivocally shown that denitrifiers can grow by respiring N_2_O ^42–44^, proving the respiratory role of N_2_OR. Moreover, an electron sink/spill role for N_2_OR has been proposed for *Gemmatimonas aurantiaca* T-27 ^45, 46^ without biomass production (i.e., growth). Despite the presence of N_2_OR in *Methylocystis* sp. SC2, its ability to grow in anoxia under N_2_O-reducing conditions is unverified ^29^. Thus, the ability to grow by converting N_2_O to N_2_ has not yet been reported for any of the known aerobic or anaerobic methanotrophs even with non-methane substrates such as methanol.

Apparent O_2_ requirements of aerobic methanotrophs contradict their high relative abundance in extremely hypoxic and even anoxic environments such as peat bogs, wetlands, rice paddies, forest soils, and geothermal habitats ^47, 48^. It is therefore critical to investigate the ability of aerobic methanotrophs with N_2_OR to use N_2_O as the sole terminal electron acceptor for energy conservation and biomass production, a metabolic trait that could allow them to thrive in these anoxic ecosystems. We used a multi-faceted approach to investigate the role of N_2_O respiration in defining the physiology and ecology of selected aerobic methanotrophs. Growth experiments demonstrated that the presence of N_2_OR in an acidophilic proteobacterial methanotroph, *Methylocella tundra* T4, and an extremely acidophilic verrucomicrobial methanotroph, *Methylacidiphilum caldifontis* IT6, enables these organisms to respire N_2_O and to produce biomass while oxidizing a wide variety of electron donors, including methanol, acetol, pyruvate, and hydrogen. Compared to N_2_O, respiration of NO_3_^-^ and NO_2_^-^ did not support the growth of these methanotrophs on C1 substrates in anoxic environments. We also demonstrate that *Methylocella tundra* T4 can reduce both O_2_ and N_2_O simultaneously, allowing it to oxidize more CH_4_ and generate more biomass under O_2_-limiting conditions. Our findings significantly expand the potential ecological niche of aerobic methanotrophs and reveal that some methanotrophic microbial strains could be used for mitigating multiple GHGs.

## Results and discussion

### N_2_OR-encoding genes in aerobic methanotrophs

To identify methanotrophs capable of using N_2_O as an alternative electron acceptor, publicly available genomes, and metagenome-assembled genomes (MAGs) of methanotrophs were screened for *nosZ* gene. We found genes encoding N_2_OR in the genomes and MAGs of three phyla containing methanotrophs: *Pseudomonadota*, *Verrucomicrobiota*, and *Gemmatimonadota* (Supplementary Table 1). They were confined to the alphaproteobacterial methanotrophs and absent in gammaproteobacterial methanotrophs in the case of the phyla *Pseudomonadota* and represented only by two genera, *Methylocella* and *Methylocystis*, which also accounted for the majority of the methanotrophs genomes encoding *nosZ*. Similarly, *nosZ* genes were exclusively found in a representative genome in the phyla *Verrucomicrobiota* (represented by the genus *Methylacidiphilum*) and *Gemmatimonadota* (represented by the candidate genus ‘Methylotropicum’), respectively. Phylogenetic analysis of predicted NosZ protein sequences revealed that the majority of N_2_OR-containing methanotrophs belong to clade I. The nosZ genes found in *Methylocella* and *Methylocystis* are from the clade I NosZ lineage, while those found in *Methylacidiphilum* and ‘*Ca.* Methylotropicum’ are from the clade II NosZ lineage (Fig. 1 and Supplementary Fig. 1).

**Fig. 1.**
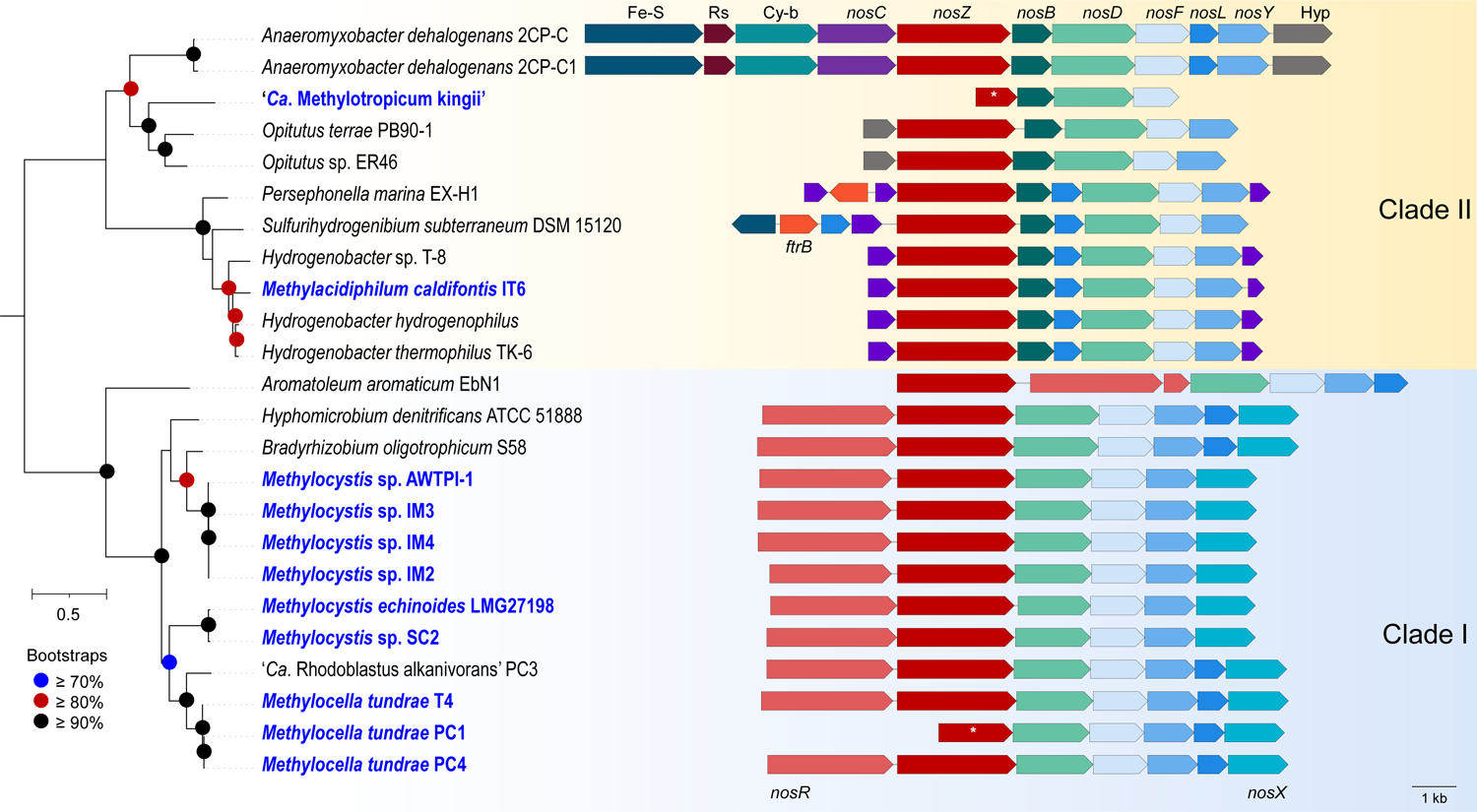
A maximum-likelihood phylogenetic tree of NosZ proteins, with *nos* operon arrangements in methanotrophic and non-methanotrophic bacterial strains. The phylogenetic tree was constructed with IQ-TREE (IQ-TREE options: -B 1000 -m LG+F+R5) using aligned NosZ (details in *Materials and Methods*) and rooted at the mid-point. The scale bar represents a 0.5 change per amino acid position. Organization of the *nos* operon in methanotrophic strains (labeled in blue text) and closely related non-methanotrophic bacteria are shown. The genes, represented by arrows, are drawn to scale. Homologs are depicted in identical colors. The NosZ amino acid sequences and gene arrangement information were retrieved using the following genome accessions: GCF_017310505.1, *Methylacidiphilum caldifontis* IT6; GCF_000010785.1, *Hydrogenobacter thermophilus* TK-6; GCF_011006175.1, *Hydrogenobacter* sp. T-8; GCF_900215655.1, *Hydrogenobacter hydrogenophilus* DSM 2913; GCF_000619805.1, *Sulfurihydrogenibium subterraneum* DSM 15120; GCF_000021565.1, *Persephonella marina* EX-H1; GCF_000022145.1, *Anaeromyxobacter dehalogenans* 2CP1; GCF_000013385.1, *Anaeromyxobacter dehalogenans* 2CP-C; GCF_003054705.1, *Opitutus* sp. ER46; GCF_000019965.1, *Opitutus terrae* PB90-1; GCF_901905185.1, *Methylocella tundrae* PC4; GCA_901905175.1, *Methylocella tundrae* PC1; CP139089.1, *Methylocella tundrae* T4; FO000002.1, *Methylocystis* sp. SC2; GCF_000025965.1, *Aromatoleum aromaticum* EbN1; GCF_022760775.1, ‘*Candidatus* Rhodoblastus alkanivorans’ PC3; GCF_000143145.1, Hyphomicrobium denitrificans ATCC 51888; GCF_000344805.1, *Bradyrhizobium oligotrophicum* S58; GCF_027923385.1, *Methylocystis echinoides* LMG27198; GCA_003963405.1, *Methylocystis* sp. AWTPI-1. * indicates that the *nosZ* genes are truncated due to genome fragmentation. Source Data contains annotation information for *Methylocystis* spp. IM2, IM3, and IM4, as well as ‘*Ca.* Methylotropicum kingii’.

Three *Methylocella tundrae* strains: T4 (re-sequenced genome), PC1 ^49^, and PC4 ^49^, have *nos* gene clusters (NGC) (Fig. 1). These are incorporated into *nosRZDFYLX* operons in strains PC4 and T4 and a *nosZDFYLX* operon in strain PC1 (Fig. 1). Strain PC1 has truncated *nosZ* and missing *nosR* genes. This is most likely due to its genome being highly fragmented into several small contigs containing missing and fragmented genes. The NGC composition and operon arrangement, n*osRZDFYLX*, were largely similar in the genomes of the six N_2_OR-containing *Methylocystis* species (Fig. 1), including *Methylocystis* sp. SC2 ^39^, *Methylocystis echinoides* LMG27198, three in-house *Methylocystis echinoides*-like isolates (strains IM2, IM3, and IM4), and a metagenome-assembled genome (MAG) of a *Methylocystis* sp. AWTPI-1 recovered from a water treatment facility ^50^. A notable feature in their NGC organization was the absence of the gene encoding the membrane-anchored copper chaperon, NosL, which is primarily involved in Cu(I) delivery to apo-NosZ ^51, 52^. Methanotrophs with copper-containing MMO usually possess multiple copper chaperones ^53, 54^ that may complement NosL, making it non-essential for NosZ maturation. Altogether, the NGC in these alphaproteobacterial methanotrophs has a similar organization to those of clade I N_2_O-reducers (Fig 1). BLAST results further revealed that the individual *nos* genes in the *Methylocella* and *Methylocystis* strains shared a high degree of similarity to each other and other non-methanotrophic *Alphaproteobacteria* (Supplementary Table 2). Also, their NosZ proteins share high homology with proteins annotated as twin-arginine translocation (Tat)-dependent N_2_OR (35–89%) and also possess the Tat signal peptide with a characteristic SRRx[F|L] motif ^55^ found in clade I NosZ ^38^.

The NGC in the genome of *Methylacidiphilum caldifontis* IT6 ^56^, comprises a *nosCZBLDFYC* operon (Fig. 1) but lacks the typical *nosX* and *nosR* found in clade I N_2_O-reducers ^37, 38^, involved in NosR maturation ^57, 58^ and electron transfer to NosZ ^59, 60^, respectively. Notably, the NGC (IT6_00904–00911) was found within the cluster of genes (IT6_00903, IT6_00912–00917) encoding Alternative Complex III. Both the *aa*3-type and *cbb*3-type cytochrome *c* oxidase-encoding genes are also located right next to these genes. Genes encoding two *c*-type cytochromes (*nosC*) within the *nos* operon (Fig. 1) could serve electron transport functions ^61^. Interestingly, BLAST and synteny analyses of the NGC show that the individual genes are most closely related to similar genes found in genomes of an extremely thermophilic *Hydrogenobacter* species of the phylum *Aquificota* (amino acid identities of 72.41–91.96%) (Supplementary Table 2) with a similar genetic organization (Fig. 1). Strain IT6 NosZ shares high similarities to proteins annotated as Sec-dependent N_2_OR (35–89%) with an N-terminal Sec-type signal peptide found in clade II NosZ ^37, 38^; the highest identities (79–89%) were with NosZ proteins from other *Hydrogenobacter* species. *Hydrogenobacter thermophilus* TK-6, a hydrogen-oxidizing bacterium, can completely denitrify NO_3_ ^-^ to N gas ^62^, indicating the presence of a functional N_2_OR. As a result, *Methylacidiphilum caldifontis* IT6 may also have a functional N_2_OR due to the high similarity of its NGC to those of *Hydrogenobacter* species. Although genomes of other *Methylacidiphilum* species, including *Methylacidiphilum fumariolicum,* were observed to lack the gene encoding the N_2_OR catalytic subunit, NosZ, some genes encoding Nos accessory proteins were found (Supplementary Table 2). Interestingly, the N_2_OR genes for *Methylacidiphilum caldifontis* IT6 were found in a genomic island (Supplementary Table 3) and were most likely acquired through horizontal gene transfer, which is consistent with its NosZ phylogeny (Fig. 1, Supplementary Fig. 1). This is not surprising since many key metabolic genes in verrucomicrobial methanotrophs, including those encoding the methane monooxygenases, are believed to have been acquired through horizontal gene transfer ^63^. As a result, *Methylacidiphilum fumariolicum* strains might have likely acquired the NGC before losing the key functional genes but retaining some of the accessory genes. Finally, we found a *nosZBDF* operon in the MAG of the uncultured methanotrophic bacterium ‘*Ca*. Methylotropicum kingii’ ^64^ that resembles clade II NGC, with a truncated *nosZ* and multiple missing genes like *nosY*, *nosL*, and *nosC* (Fig. 1). These are also likely the result of multiple MAG fragmentations.

Multiple sequence alignments of the predicted NosZ proteins of methanotrophs and other microorganisms (clade I and II) were constructed. All the expected metal-binding residues present in N_2_OR were mostly conserved in the methanotroph NosZ sequences (Supplementary Fig. 2). The nine histidine residues of the beta-propeller domain implicated in Cu_Z_-binding were completely conserved. Like the clade I NosZ proteins, the Cu_Z_-binding motifs of the *Methylocystis* and *Methylocella* strains associated with the first two histidine (DxHHxH) and the last histidine (EPHD), were also completely conserved. These residues were less completely conserved in strain IT6 (i.e., DxHH and EPH) as seen in other clade II NosZ proteins. Considering the extremely acidophilic and thermophilic nature of strain IT6, the conservation of the key residues is consistent with its function as a NosZ protein.

### N_2_O-dependent anaerobic growth of methanotrophs

The presence of genes predicted to encode N_2_OR in the genomes of *Methylocella tundrae* strains, *Methylacidiphilum caldifontis* IT6, and *Methylocystis* strains (SC2, IM2, IM3, and IM4) (Supplementary Table 1) led us to investigate whether this enzyme can support the anaerobic growth of these aerobic methanotrophs when N_2_O is supplied as their sole electron acceptor. Physiological studies on N_2_O reduction by methanotrophs focused on *Methylocella tundrae* T4 and *Methylacidiphilum caldifontis* IT6 since preliminary experiments showed that the N_2_OR-containing *Methylocystis* strains, including *Methylocystis* sp. SC2 and the in-house *Methylocystis* strains (IM2, IM3, and IM4) failed to reduce N_2_O under various anoxic growth conditions. We set up anoxic batch cultures of *Methylocella tundrae* T4 and *Methylacidiphilum caldifontis* IT6 using methanol as a sole electron donor with or without N_2_O as the sole electron acceptor. For these incubations, 2 mM ammonia (NH_4_ ^+^) was used as the nitrogen source instead of NO_3_ ^−^ to avoid the involvement of dissimilatory nitrate reduction particularly in the *Methylocella* strains with nitrate-reducing potential. As a negative control, closely related methanotrophs lacking a predicted N_2_OR (*Methylocella silvestris* BL2 and *Methylacidiphilum infernorum* IT5, respectively) were included in the study design. The growth experiments were conducted in LSM medium at pH 5.5 for *Methylocella* species (strains T4 and BL2) and at pH 2.0 for *Methylacidiphilum* species (strains IT5 and IT6). As expected, in control incubations provided with O_2_ as terminal electron acceptor, all four strains grew on CH_3_OH (Figs. 2a, 2d, 2g, 2j). In these controls, *Methylocella* strains displayed higher growth rates (strain T4: *µ*_max_ = 2.83 ± 0.03 d^−1^; strain BL2: *µ*_max_ = 1.79 ± 0.05 d^−1^) than *Methylacidiphilum* strains (strain IT6: *µ*_max_ = 1.57 ± 0.04 d^−1^; strain IT5: *µ*_max_ = 1.49 ± 0.01 d^−1^).

**Fig. 2.**
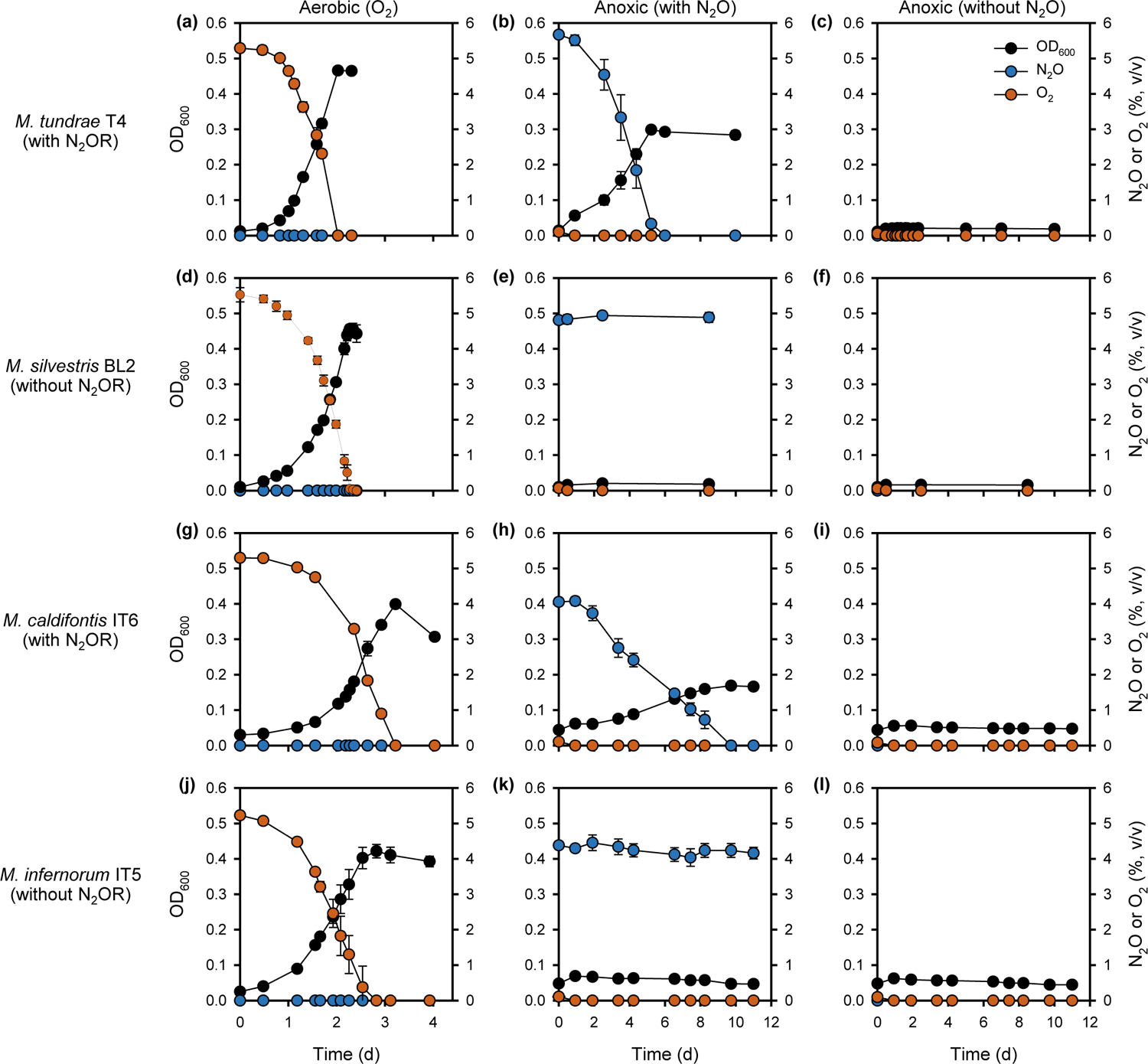
Aerobic and anaerobic growth of N_2_OR-containing and N_2_OR-lacking *Methylocella* and *Methylacidiphilum* strains on methanol. *Methylocella tundrae* T4, *Methylocella silvestris* BL2, *Methylacidiphilum caldifontis* IT6, and *Methylacidiphilum infernorum* IT5 cells were grown in LSM medium supplemented with 30 mM methanol as electron donor and NH_4_ ^+^ as N-source. Aerobic growth of the 4 strains with O_2_ (**a**, **d**, **g**, and **j**), anaerobic growth with N_2_O (**b**, **e**, **h**, and **k**), and anaerobic growth without N_2_O (**c**, **f**, **i**, and **l**) as the sole terminal electron acceptor was determined by optical density measurements at 600 nm, followed by measurements of O_2_ and N_2_O consumption in the headspace of the culture bottles. Note that the trace O_2_ present at the start of the incubation in the anaerobic cultures without N_2_O did not contribute to obvious growth (**c**, **f**, **i**, and **l**). All experiments were performed in triplicates. Data are presented as meanD±D1 SD, and the error bars are hidden when they are smaller than the width of the symbols.

Under N_2_O-containing anoxic conditions, *Methylocella tundrae* T4 and *Methylacidiphilum caldifontis* IT6 reduced N_2_O and grew anoxically on methanol (Figs. 2b, 2h). When N_2_O was depleted, the growth of strains T4 and IT6 ceased. No growth was observed in N_2_O-free anoxic conditions used as negative controls (Figs. 2c, 2i). These results demonstrate that the anoxic growth of these methanotrophs was dependent on N_2_O as the sole electron acceptor. The observed N_2_O reduction was catalyzed by a functional respiratory N_2_OR, as the N_2_OR-lacking relatives (*Methylacidiphilum infernorum* IT5 and *Methylocella silvestris* BL2) used as negative controls did not grow or reduce N_2_O under anoxic conditions (Figs. 2e, 2f, 2k, 2l). In addition, other known electron donors of *Methylocella tundrae* T4 and *Methylacidiphilum caldifontis* IT6, which support their aerobic growth ^56, 65, 66^, also supported their growth under anoxic N_2_O-reducing conditions (Supplementary Table 4). *Methylocella tundrae* T4 grew on pyruvate and acetol, while *Methylacidiphilum caldifontis* IT6 grew on acetol under anoxic N_2_O-reducing conditions. Further, molecular hydrogen supported the chemolithoautotrophic growth of *Methylacidiphilum caldifontis* IT6 as the sole electron donor under anoxic N_2_O-reducing conditions (Supplementary Fig. 3). The transcriptomic analysis (see below) suggests that the group 1d [NiFe] hydrogenase encoded in the genome of *Methylacidiphilum caldifontis* IT6 could be involved in chemolithoautotrophic growth under anoxic N_2_O respiring conditions.

*Methylocella tundrae* T4 exhibited a higher growth rate (*µ*_max_ = 0.47 ± 0.02 d^−1^) than *Methylacidiphilum caldifontis* IT6 (*µ*_max_ = 0.18 ± 0.01 d^−1^) on methanol and N_2_O. However, these values are approximately 6 and 9 times, respectively, lower than the growth rates measured for both strains under O_2_−respiring conditions. Biomass yields Y*_x/m_* (g DW⋅mol^−1^ N_2_O or O_2_ reduced) for the methanol-oxidizing cultures of strains T4 and IT6 reducing N_2_O as the sole electron acceptor were also lower than for cells reducing O_2_ as the sole electron acceptor. The biomass yield of *Methylocella tundrae* T4 cells grown anaerobically on N_2_O (4.64 ± 0.04 g DW⋅mol^−1^ N_2_O reduced) was approximately 45% of that of aerobically grown cells (10.41 ± 0.04 g DW⋅mol^−1^ O_2_ reduced). Similarly, *Methylacidiphilum caldifontis* IT6 had a biomass yield when grown anoxically on N_2_O (2.36 ± 0.04 g DW⋅mol^−1^ N_2_O reduced), which was only about 38% of that achieved by aerobically grown cells (6.27 ± 0.14 g DW⋅mol^−1^ O_2_ reduced). This improved molar yield on O_2_ is expected despite the higher reduction potential of N_2_O (see Eqs. [1] and [2]), since O_2_ respiration accepts twice as many electrons as N_2_O respiration (Eqs. 1 & 2) ^67^. In addition, the terminal oxidases of both strains are proton pumps and conserve energy (Supplementary Tables 5, 6) ^68, 69^, whereas N_2_OR does neither ^70^. To our knowledge, our results constitute the first report of N_2_O reduction coupled with anaerobic growth in any methanotroph.

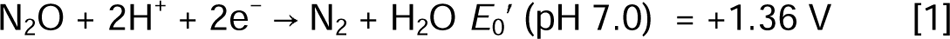

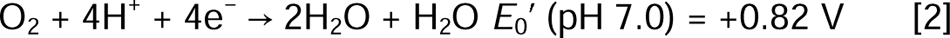

It is well known that N_2_O reduction is generally inhibited at acidic pH (< 6.0) ^71, 72^, resulting in N_2_O accumulation in acidic environments ^33, 73^. However, the current study revealed that two acidophilic methanotrophs, *Methylocella tundrae* T4 and *Methylacidiphilum caldifontis* IT6 can reduce N_2_O in moderately acidic (pH 5.5) and extremely acidic (pH 2.0) conditions, respectively. The existence of acid-tolerant N_2_O reducers (pH 4.0 to 6.0) has been proposed in soil microcosm and enrichment experiments ^74, 75^. So far, the only isolate implicated in N_2_O reduction at an acidic pH (5.7) is a *Rhodanobacter* species (strain C01) isolated from acidic soil in Norway ^76^. Our study reveals that N_2_O reduction can occur even at an extremely acidic pH of 2.0. Furthermore, the conditions required for N_2_O reduction in the N_2_OR-containing *Methylocystis* strains remain unresolved. Perhaps some unknown growth or environmental factors are required to stimulate N_2_O respiration in these methanotrophs, which will require further investigation.

### Nitrate and nitrite reduction in Methylocella spp

#### No anoxic growth of Methylocella spp. with CH_3_OH and NO_3_ ^-^

We next tested if the presence of denitrification enzymes in *Methylocella tundrae* T4 (nitrate reductase [NAR], nitric oxide reductase [NOR] and N_2_OR) and *Methylocella silvestris* BL2 (NAR, nitrite reductase [NIR], and NOR) (Supplementary Table 1) can equate to growth when NO_3_ ^−^ or NO_2_ ^−^ is used as the sole terminal electron acceptor. Indeed, the presence of NAR (and NIR) in these methanotrophs resulted in NO_3_ ^−^ (and NO_2_ ^−^) reduction when methanol was provided as the sole electron donor, however, growth was barely detected under these conditions (Fig. 3a, 3b). Strain T4, which lacks a canonical NIR, reduced all the provided NO_3_ ^−^ stoichiometrically to NO_2_ ^−^ when provided with methanol as the sole electron donor (Fig. 3a). Under the same condition, strain BL2, a NAR and NIR-containing methanotroph, initially reduced the provided NO_3_ ^−^ to NO_2_ ^−^ and eventually, all the accumulated NO_2_ ^−^ was stoichiometrically reduced to N_2_ O towards the end of the incubation (Fig. 3b). These results demonstrate that these methanotrophs have a functional NAR and or NIR and can utilize NO_2_ ^−^ and/or NO_2_ ^−^ instead of O_2_ as a terminal electron acceptor. Nevertheless, these methanotrophs do not appear to rely on these activities for growth. Likewise, other aerobic methanotrophs have demonstrated denitrification activities under suboxic conditions. For example, the gammaproteobacterial methanotrophs *Methylomonas denitrificans* FJG1 and *Methylomicrobium album* BG8 were discovered to couple the oxidation of diverse electron donors to NO_3_ ^−^ and NO_2_ ^−^ reduction, respectively ^26, 77^. However, none of these strains was demonstrated to couple this activity to growth. It should be noted that the genomes of all known *Methylacidiphilum* strains lack genes encoding a respiratory NAR (Supplementary Table 1).

**Fig. 3.**
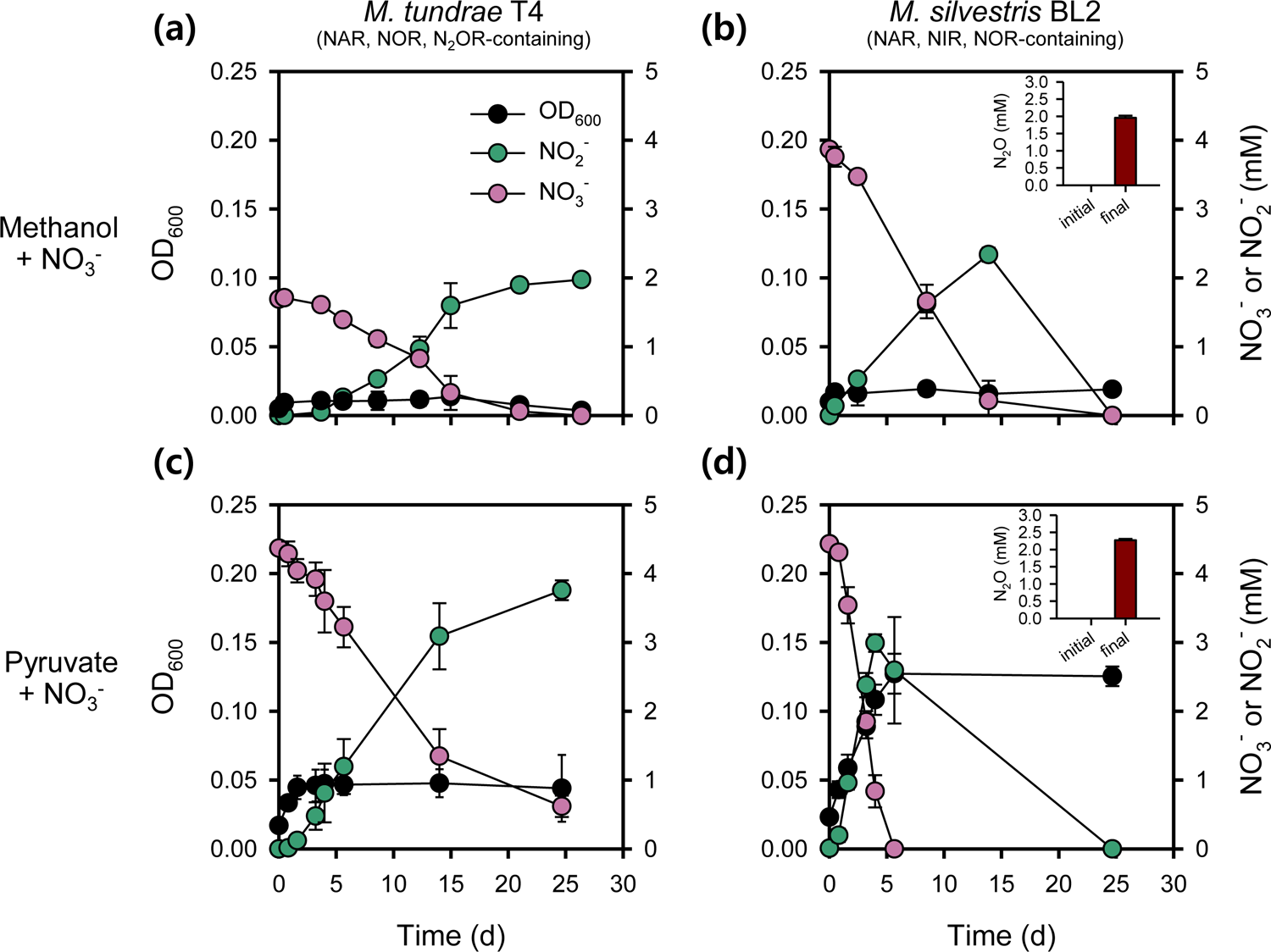
Anaerobic growth of *Methylocella* strains on methanol or pyruvate as the sole electron donor and NO ^−^ as the terminal electron acceptor. *Methylocella tundrae* T4 and *Methylocella silvestris* BL2 cells were grown in LSM medium supplemented with 30 mM methanol and 2–4 mM NO_3_ ^−^. NH_4_ ^+^ (2 mM) was supplied as the N-source. Anaerobic growth of *Methylocella tundrae* T4 (**a**) and *Methylocella silvestris* BL2 (**b**) cells on methanol as the sole electron donor with NO_3_^−^ as the sole electron acceptor. Anaerobic growth of *Methylocella tundrae* T4 (**c**) and *Methylocella silvestris* BL2 (**d**) cells on pyruvate as the sole electron donor with NO_3_^−^ as the sole electron acceptor. N_2_O produced from NO_3_^−^ reduction by cells of *Methylocella silvestris* BL2 grown on methanol or pyruvate are shown as inset plots within each figure. N_2_O production was not observed in strain T4, hence inset plots for N_2_O production were not displayed. Lower NO_3_ ^−^ (ca. 2.0 mM) was used in the case of methanol (**a**) to avoid NO_2_ ^−^ toxicity. Growth was determined by optical density measurements at 600 nm, followed by measurements of NO_3_ ^-^ and NO_2_ ^−^ concentrations. Data are presented as meanD±D1 SD of triplicate experiments, and the error bars are hidden when they are smaller than the width of the symbols.

#### Toxicity of reactive nitrogen species for Methylocella spp

Considering that methanol oxidation was coupled to N_2_O reduction and led to obvious growth in the N_2_OR-containing methanotrophs (Fig. 2b, 2h), the lack of growth during NO_3_ ^−^ reduction by these microorganisms is suspected to be caused by the accumulation of growth-arresting reactive nitrogen species (RNS) metabolites like NO_2_ ^−^ and nitric oxide (NO) ^78–80^. Consistent with this hypothesis, the accumulation of NO_2_ ^−^ in suboxic cultures of *Vibrio cholerae* and other bacterial species was found to limit population expansion but nitrate reduction still promoted cell viability ^81^. NO_2_ ^−^ typically accumulates due to a lack of functional NIR as observed for strain T4 (Fig. 3a) and, to some degree, even transiently accumulates in the presence of a functional NIR, as observed for strain BL2 (Fig. 3b). The impact of NO_2_ ^−^ accumulated from NO_3_ ^−^ reduction might be more severe in acidic environments since protonation of NO_2_ ^−^ leads to the formation of free nitrous acid (FNA), a known inhibitor of microbial anabolic and catabolic processes ^82, 83^. In addition, chemodenitirification of NO ^−84, 85^ could result in an accumulation of NO in the cell environment, which is highly toxic to microbial life ^86, 87^. To further support the hypothesis of RNS toxicity, strain T4 was cultivated under N_2_O-reducing conditions with methanol as the sole electron donor and supplied with NO_3_ ^−^ instead of NH_4_ ^+^ as the N source in the medium (Supplementary Fig. 4). Consistent with the idea that NO_4_ ^−^ accumulation results in growth arrest, the culture growth plateaued at approximately the same time NO_3_ ^−^ accumulated (≥ 0.3 mM NO_2_ ^−^) (Supplementary Fig. 4a), whereas in control cultures containing NH_4_ ^+^ instead of NO_3_ ^−^ as the N-source, NO_2_ ^−^ accumulation was not observed, and the cells were able to reach higher cell densities (Supplementary Fig. 4b). Furthermore, the effect of NO_2_ ^−^ stress induced in strain T4 was verified by adding varying NO_2_ ^−^ concentrations (0, 0.01, 0.03, 0.1, 0.3, and 1 mM) to aerobic (Supplementary Fig. 5a) and anaerobic N_2_O-respiring cultures (Supplementary Fig. 5b). Nitrite, particularly at concentrations higher than 0.3 mM at pH 5.5, induced a stress in *Methylocella tundrae* T4, resulting in growth inhibition (Supplementary Fig. 5). These results are comparable to that of *Methylophaga nitratireducenticrescens* JAM1, a facultative methylotroph, which, when grown aerobically on methanol at pH 7.4, had a four-fold decrease in biomass in the presence of 0.36 mM NO_2_ ^−^ and did not grow in the presence of 0.71 mM NO_2_ ^−88^. Taken together, our data suggest that the lack of growth seen in NO_3_ ^−^/NO_2_ ^-^reducing methanotrophs when grown on methanol may be a result of RNS toxicity. On the other hand, when N_2_O is reduced to N_2_ by N_2_O-reducing methanotrophs, the creation of these RNS is avoided, which may explain the disparity in growth with N_2_O as the terminal electron acceptor compared to NO_3_ ^−^ and NO_2_ ^−^.

#### Toxicity of C1 metabolites in nitrate-reducing Methylocella spp

Aside from the inhibitory effects of RNS, toxic intermediates from methanol metabolism might synergistically contribute to the inability of methanotrophs to grow when respiring NO_3_ ^−^/NO_2_ ^−^. Although formaldehyde is a key intermediate in the C1 metabolic pathway in many methylotrophs, it is highly toxic ^89, 90^. Therefore, in situations where biomass production is limited due to RNS toxicity, it is likely that formaldehyde further retards the growth of denitrifying methanotrophs. To investigate this mechanism, we grew *Methylocella* strains under NO_3_ ^−^-reducing conditions using a C-C electron donor, pyruvate, which does not generate formaldehyde as a major metabolite (Figs. 3c, 3d). Eventually, nearly all the supplied NO_3_ ^−^ was stoichiometrically converted to NO_2_ ^−^ and N_2_ O in strains T4 and BL2, respectively. In contrast to the lack of growth on methanol, pyruvate supported the growth of both *Methylocella* strains under NO_3_ ^−^-reducing conditions (Figs. 3c, 3d). Growth was more pronounced in strain BL2 than in strain T4 (Fig. 3c, 3d), possibly due to the presence of NIR and NOR in addition to NAR in strain BL2, which limited NO_2_ ^−^ accumulation (Fig. 2d). Nonetheless, no further growth on pyruvate was observed in strain BL2 after day 5, despite reduction of the accumulated NO_2_ ^−^ (∼ 2.5 mM) to N_2_O (Fig. 3d). It is worth noting that the accumulated NO_2_ ^−^ concentration is higher than the 0.3 mM concentration that inhibited *Methylocella tundrae* T4 and may also be responsible for the lack of growth in strain BL2.

Overall, these results demonstrate that in the tested *Methylocella* strains: (i) RNS have a major inhibitory effect on growth under denitrifying conditions; (ii) there are no growth benefits from methanol oxidation coupled to NO_3_ ^−^ reduction, probably due to toxicity of C1 metabolic intermediates as well as RNS; and (iii) anaerobic growth is observed when NO_3_ ^−^ reduction is coupled to the oxidation of pyruvate, a C-C electron donor; although the amount of growth is dependent on the completeness of the denitrification pathway and the accumulation of RNS. These propositions are supported by increased expression of genes involved in RNS and C1 metabolite detoxification under denitrifying conditions (see transcriptomic analysis below). Taken together, these results may explain why methanotrophs that couple methanol oxidation to NO_3_ ^−^ or NO_2_ ^−^ reduction show no clear signs of growth due to this process. Most methanotrophs can only utilize methane and its C1 derivatives as energy sources ^16, 91^ and thus should not be able to grow under denitrifying conditions ^26, 27^. On the other hand, versatile facultative methanotrophs of the genus *Methylocella* are potentially able to grow in strictly anoxic habitats when C-C bond substrates are available. In natural systems, RNS toxicity to methanotrophs can be lowered due to low *in situ* NO_3_ ^−^/ NO_2_ ^−^ concentrations and/or coexistence with other NO_2_ ^−^-reducing microorganisms, implicating possible growth of methanotrophs by anoxic NO_3_ ^−^/NO_2_ ^−^ reduction.

### N_2_O reduction coupled with CH_4_ oxidation

#### N_2_O reduction kinetics

We investigated N_2_O respiration kinetics using resting cells of anaerobic N_2_O-respiring cultures (CH_3_OH + N_2_O) in a microrespirometry (MR) chamber. Harvested cells of strains *Methylacidiphilum caldifontis* IT6 and *Methylocella tundrae* T4 were dispensed into a closed 10-mL MR chamber outfitted with O_2_ and N_2_O-detecting microsensors, supplied with CH_3_OH (2 mM) and N_2_O as a sole electron donor, and acceptor, respectively, and incubated anoxically. The N_2_O respiration kinetics followed Michaelis-Menten kinetics (see Supplementary results, Supplementary Fig. 6). The cells of strains T4 and IT6 grown at anoxic CH_3_OH + N_2_O conditions reduced N_2_O at a maximum rate of 1.122 ± 0.005 mmol N_2_O·h^−1^·g DW^−1^ (Supplementary Fig. 6a) and 0.414 ± 0.003 mmol N_2_O·h^−1^·g DW^−1^ (Supplementary Fig. 6b), respectively. The molar ratios of CH_3_OH to O_2_ and CH_3_OH to N_2_O consumed were approximately 1:1.0 (± 0.05; *n* = 3) and 1:2.04 (± 0.17; *n* = 3), respectively, which coincide with the theoretical values obtained from the equations 3 and 4.

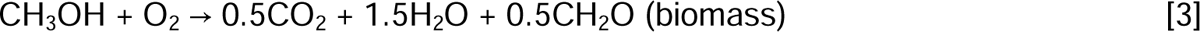

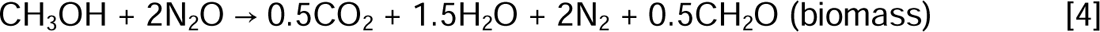

#### Sensitivity of N_2_OR to O_2_

While O_2_ is well known to impair N_2_OR activity ^34, 92^, some bacterial strains have been reported to reduce N_2_O in the presence of O_2_ ^93, 94^. We therefore tested the capacity of strains IT6 and T4 to reduce N_2_O in the presence of O_2_ by using resting cells of anoxic CH_3_OH + N_2_O-cultures. After spiking O_2_ to strain IT6 cells respiring N_2_O in the anoxic MR chamber, N_2_O-respiration ceased: dropping from the maximum (0.4–0.5 mmol N_2_O·h^−1^·g DW^−1^) to zero (Fig. 4a, Table 1). N_2_O reduction activity only started when the O_2_ concentration was below ca. 3 µM, suggesting the N_2_O reduction activity of this strain is highly sensitive to O_2_. In contrast, when O_2_ (∼ 14 and 35 µM) was added to N_2_O-respiring cells of strain T4, simultaneous reduction of N_2_O and O_2_ was observed (Fig. 4b). However, the N_2_O respiration rates dropped to 0.24 and 0.13 mmol N_2_O·h^−1^·g DW^−1^ after spiking ∼14 and 30 µM O_2_, respectively, which are approximately 34 and 20% of the maximum rate before O_2_ introduction (0.64–0.68 mmol N_2_O·h^−1^·g DW^−1^). These results suggest that N_2_O reduction in strain T4 is not highly impaired by O_2_ in comparison to strain IT6 and N_2_OR activity fully recovered in both strains as O_2_ was being depleted. Because the N_2_OR of strain IT6 was found to be highly sensitive to O_2_, further characterization of methanotroph N_2_OR activity in response to O_2_ exposure was limited to strain T4.

**Fig. 4.**
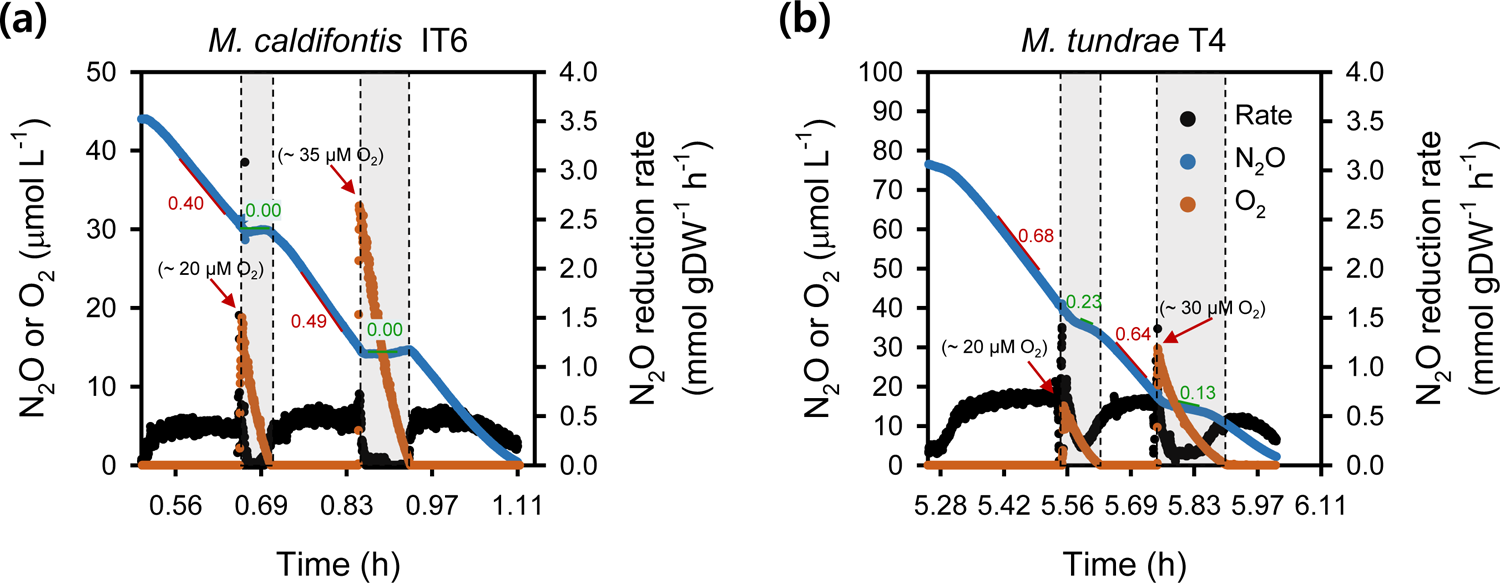
Microrespirometry-based N_2_O and O_2_ reduction by cells of *Methylacidiphilum caldifontis* IT6 (a) and *Methylocella tundrae* T4 (**b**) during methanol oxidation. Filled blue dots represent dissolved N_2_O, filled orange dots represent dissolved O_2_, and filled black dots represent N_2_O reduction rates. Experiments were performed in a microrespirometry (MR) chamber fitted with O_2_ and N_2_O microsensors. The red arrows mark the addition of 20–35DµM O_2_ into the MR chamber. The red- and green-marked numbers close to the red and green lines represent the N_2_O reduction rates before and during O_2_ reduction (gray-shaded area) in the MR chamber, respectively.

**Table 1:** Microrespirometry-based substrate-specific N_2_O- or O_2_-reduction rate by *Methylocella tundrae* T4 cells grown under anoxic and suboxic growth conditions.

| Condition | Rate (mmol·h<sup>-1</sup>·g DW<sup>-1</sup>) |
| --- | --- |
| Maximum respiration rates of anoxic CH<sub>3</sub>OH + N<sub>2</sub>O-respiring cells |  |
| N<sub>2</sub>O respiration (at 0 μM O<sub>2</sub>; electron donor CH<sub>3</sub>OH; at the first 4 spikes of N<sub>2</sub>O) | 1.12 ± 0.01 |
| N<sub>2</sub>O respiration (at 0–5 μM O<sub>2</sub>; electron donor CH<sub>3</sub>OH; at the 5<sup>th</sup> spikes of N<sub>2</sub>O) | 0.64–0.68 |
| N<sub>2</sub>O respiration (at O<sub>2</sub> > 5 μM; electron donor CH<sub>3</sub>OH) | 0.13–0.24 |
| O<sub>2</sub> respiration (at O<sub>2</sub> > 5 μM; electron donor CH<sub>3</sub>OH) | 1.02–1.07 |
| Maximum respiration rates of suboxic CH<sub>4</sub> + N<sub>2</sub>O + O<sub>2</sub>-respiring cells |  |
| N<sub>2</sub>O respiration (at 25–60 μM O<sub>2</sub>; electron donor = CH<sub>4</sub>) | 1.58–2.47 |
| N<sub>2</sub>O respiration (at 5–170 μM O<sub>2</sub>; electron donor = CH<sub>4</sub>) | 1.32 ± 0.25 |
| O<sub>2</sub> respiration (at 25–60 μM O<sub>2</sub>; electron donor = CH<sub>4</sub>) | 0.98–2.37 |
| O<sub>2</sub> respiration (at 5–170 μM O<sub>2</sub>; electron donor = CH<sub>4</sub>) | 0.95 ± 0.09 |

These values were obtained from respiration activities with cells that had CH<sub>3</sub>OH or CH<sub>4</sub> as the sole electron donor.

Considering these results, we set out to see if cells of strain T4 could continue N_2_O respiration while using O_2_ for CH_4_ oxidation in the MR chamber. The cells used for this experiment were cultured in suboxic conditions with starting gas mixing ratios (v/v) of 1% O_2_, 5% N_2_O, and 20% CH_4_ (i.e., CH_4_ + O_2_ + N_2_O condition). Similar to the anoxic CH_3_OH + N_2_O-adapted cells described above, the suboxic CH_4_ + O_2_ + N_2_O-adapted cells co-respired O_2_ and N_2_O after injecting CH_4_ (∼406 µM) into a 5-mL MR chamber containing O_2_ (∼30 µM) and N_2_O (∼480 µM) (Fig. 5a). Interestingly, the maximum N_2_O respiration rates during each O_2_ spike were 1.4 to 2 times higher (1.58–2.47 mmol N_2_O·h^−1^·g DW^−1^) in the suboxic CH_4_ + O_2_ + N_2_O-adapted cells (Fig. 5b, Table 1) than in the anoxic CH_3_OH + N_2_O-adapted cells (1.12 ± 0.01 mmol N_2_O·h^−1^·g DW^−1^) (Table 1, Supplementary Fig. 6b), suggesting that the cells can modulate the rates of N_2_O reduction in response to O_2_ availability.

**Fig. 5.**
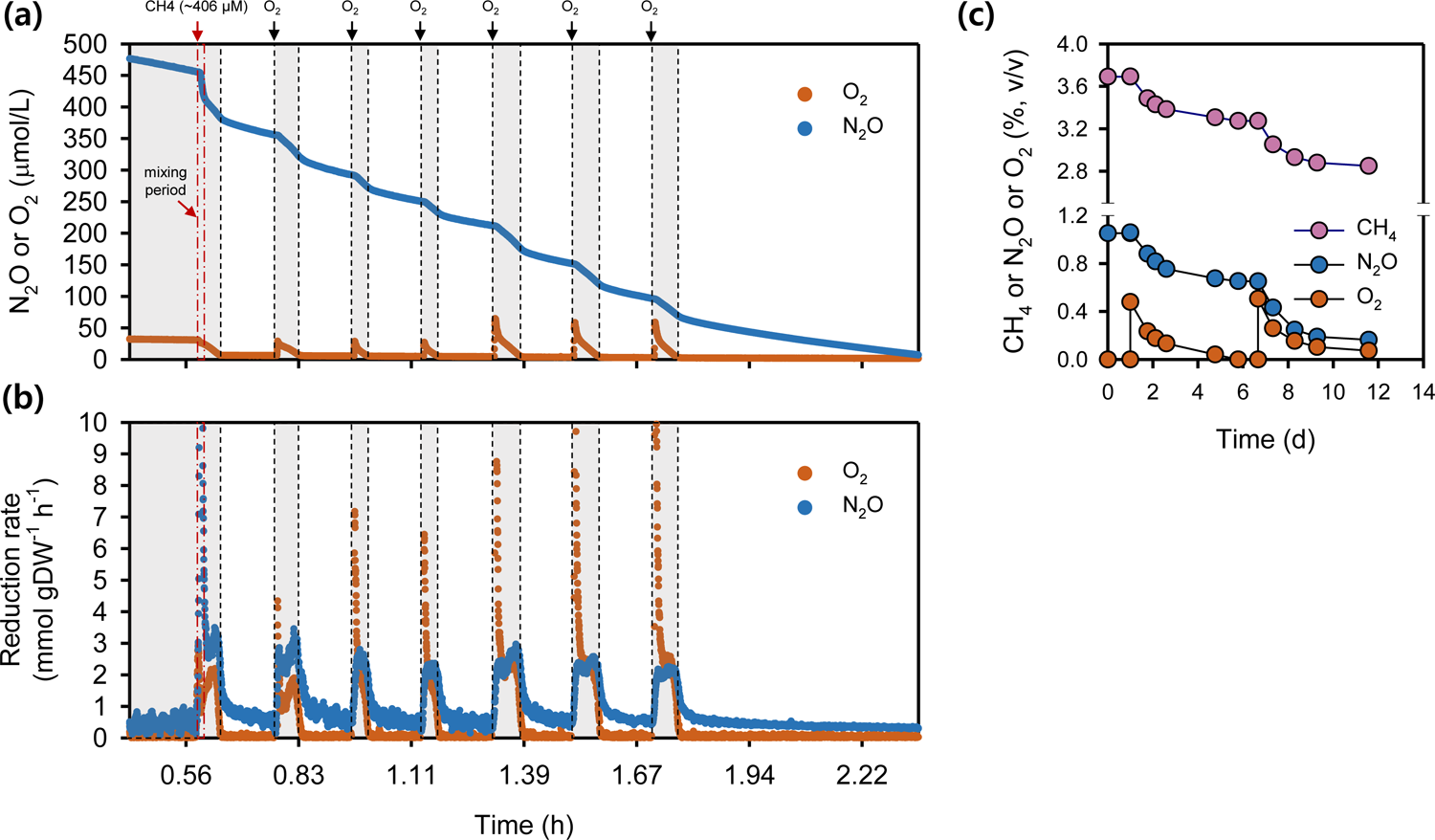
Simultaneous N_2_O and O_2_ reduction by *Methylocella tundrae* T4 cells during CH_4_ oxidation in microrespirometry (MR) and growth experiments. (**a**) MR experiment showing the simultaneous reduction of N_2_O and O_2_ by *Methylocella tundrae* T4 cells during CH_4_ oxidation. (**b**) N_2_O and O_2_ reduction rates by cells of strain T4 during CH_4_ oxidation calculated from (**a**). The filled orange and blue dots in the upper panel (**a**) represent the concentrations of dissolved O_2_ and N_2_O, respectively. The filled orange and blue dots in the bottom panel (**b**) represent the rates of O_2_ and N_2_O reduction, respectively. Experiments were performed in a MR chamber fitted with O_2_ and N_2_O microsensors. The red arrow marks the addition of CH_4_ (∼ 406 µM) into the MR chamber. Black arrow marks the addition of ∼D26 µM or ∼D60 µM O_2_ into the MR chamber. The gray-shaded area represents points where N_2_O and O_2_ are reduced simultaneously. (**c**) Growth experiment showing *Methylocella tundrae* T4 cells reducing N_2_O and O_2_ simultaneously during CH_4_ oxidation. The culture was grown in 2-liter sealed bottles (triplicates) containing 60 mL of LSM medium with 2 mM NH_4_ ^+^ as the N-source. The headspace of the bottles was composed of CH_4_ (5%, v/v), O_2_ (0.5%, v/v), N_2_O (1.4%, v/v), and CO_2_ (5%, v/v) and supplemented with additional O_2_ (∼0.5%, v/v) before its depletion. The incubation period shown in panel (**c)** is after the initial 20-day incubation period. After the depletion of O_2_, additional O_2_ was spiked to observe the simultaneous reduction of O_2_ and N_2_O during CH_4_ oxidation. Data are presented as the meanD±D1 SD of a triplicate experiment, and the error bars are hidden when they are smaller than the width of the symbols.

Accordingly, the maximum rates of N_2_O-(1.58–2.47 mmol N_2_O·h^−1^·g DW^−1^) and O_2_-reduction (0.98–2.37 mmol O_2_·h^−1^·g DW^−1^) by the suboxic CH_4_ + O_2_ + N_2_O-adapted cells were comparable (Fig. 5b, Table 1). As the O_2_ concentration and reduction rate decreased, the N_2_O reduction rate also decreased (Figs. 5a, 5b), revealing that activation of CH_4_ by O_2_ is required for stimulating N_2_O respiration by CH_4_ + O_2_ + N_2_O-adapted cells. Based on these results, we conclude that, under suboxic conditions, both aerobic CH_4_ oxidation and N_2_O reduction were operating in concert: O_2_ was needed for the monooxygenase, but the N_2_OR remained active and was able to accept electrons released downstream in the C1 oxidation pathway. This adds to the evidence that aerobic N_2_O respiration occurs in strain T4 and is linked to aerobic CH_4_ oxidation.

Finally, we attempted to ascertain the O_2_ concentration at which the suboxic CH_4_ + O_2_ + N_2_O-adapted cells of strain T4 show their N_2_O-reducing activity. At a dissolved O_2_ (DO) concentration of 170 µM, O_2_ and N_2_O were reduced simultaneously (Supplementary Figs. 7a, 7b). The maximum N_2_O reduction rate (Table 1) was nearly constant (1.32 ± 0.25 mmol N_2_O·h^−1^·g DW^−1^) across the DO range of 5–170 µM (Supplementary Figs. 7b, 7c) and was about 1.4 times higher than the maximum O_2_ reduction rates (0.95 ± 0.09 mmol O_2_·h^−1^·g DW^−1^). This means that even when exposed to high levels of O_2_, the N_2_OR in the suboxic CH_4_ + O_2_ + N_2_O-adapted cells remained functional and could reduce N_2_O at high rates. Other bacterial strains’ N_2_OR activities have been reported at DO concentrations between 100 and 260 µM ^93, 94^, indicating that their N_2_OR activity is similarly O_2_-tolerant ^93^ as that of strain T4. According to the findings of Wang and colleagues^93^, N_2_O reducers with an O_2_ tolerant N_2_OR maintain low internal O_2_ concentrations in their cells by rapidly consuming O_2_, allowing the N_2_OR to remain active. However, it remains unclear if *Methylocella tundrae* T4 also employs a similar strategy to maintain an O_2_-tolerant N_2_OR.

#### Improved methanotrophic growth of Methylocella tundrae in the presence of N_2_O

Based on the MR experiments showing the simultaneous reduction of O_2_ and N_2_O by CH_4_-fed cells of strain T4, alongside the clear N_2_O-dependent anaerobic growth, we hypothesized that strain T4 growth can be enhanced when it oxidizes CH_4_ by simultaneously reducing O_2_ and N_2_O under suboxic conditions. Using fed-batch growth experiments, we verified that strain T4 grows by CH_4_ oxidation coupled with co-respiration of N_2_O and O_2_ (Fig. 5c, Table 2), strongly supporting the MR results above. Cells grown under the suboxic CH_4_ + O_2_ + N_2_O condition consumed roughly the same amount of O_2_ and N_2_O (Fig. 5c, Table 2), and these values were comparable to what CH_4_ + O_2_ + N_2_O-grown cells consumed in the MR experiments (see Fig. 5a). Consequently, our results demonstrate that in an O_2_-limited environment, the cells can benefit energetically by directing more O_2_ to the monooxygenase step of CH_4_ oxidation, and simultaneously running a hybrid (O_2_ + N_2_O) electron transport system as shown in Table 2 and Fig. 5c.

**Table 2:** The effect of N_2_O addition on CH_4_-oxidizing cultures of *Methylocella tundrae* T4 growing in suboxic conditions.

| Culture condition | CH<sub>4</sub> oxidized (mmol/L) | O<sub>2</sub> reduced (mmol/L) | N<sub>2</sub>O reduced (mmol/L) | Increase in OD<sub>600</sub> |
| --- | --- | --- | --- | --- |
| CH<sub>4</sub> + O<sub>2</sub> | 9.74 ± 0.39 | 14.93 ± 0.43 | 0.00 | 0.114 ± 0.006 |
| CH<sub>4</sub> + O<sub>2</sub> + N<sub>2</sub>O | 12.19 ± 0.24 | 13.96 ± 0.41 | 10.15 ± 0.35 | 0.143 ± 0.002 |

The experiment was performed in 2-liter sealed bottles (replicates) with 60 mL of LSM medium in an O<sub>2</sub>-limiting suboxic headspace with and without N<sub>2</sub>O (0.5% O<sub>2</sub>, 5% CH<sub>4</sub>, 5% CO<sub>2</sub>, and 0 or 1% N<sub>2</sub>O). Following the observation of N<sub>2</sub>O reduction in bottles containing N<sub>2</sub>O, the headspace O<sub>2</sub> and N<sub>2</sub>O mixing ratios in the bottles were increased to approximately 1% and 2% (v/v), respectively. The reduction of N<sub>2</sub>O by the cultures increased CH<sub>4</sub> oxidation and biomass compared to cultures containing only O<sub>2</sub>. Data are presented as mean ± 1 SD ( $n = 3$ )

The data showed unequivocally that the total electron equivalents released during CH_4_ oxidation to CO_2_ could account for the total electron acceptor (O_2_ + N_2_O) reduced. Based on a CH_4_ to O_2_ ratio of 1:1.57 ^112^, the total amount of O_2_ reduced (13.96 mmol/L) by the suboxic CH_4_ + O_2_ + N_2_O cultures could theoretically only account for 8.89 mmol/L oxidized CH_4_, hence an excess of 3.29 mmol/L CH_4_ was oxidized which must have required an additional electron acceptor (i.e. N_2_O). Consistently, about 10.15 mmol/L N_2_O was reduced by the suboxic CH_4_ + O_2_ + N_2_O cells, equivalent to 5.08 mmol/L O_2_, since half as many electrons are consumed per mol during N_2_O reduction to N_2_ compared to O_2_ reduction to H_2_O. By running the N_2_O respiration system, the cells lower their O_2_-demand for respiration by the terminal oxidase and maximize O_2_ use by the methane monooxygenase ^16, 95^. Due to having more CH_4_ oxidized per O_2_ reduced (∼37%) when N_2_O is present, higher cell densities (OD_600_) per O_2_ reduced (∼34%) were reached in the suboxic CH_4_ + N_2_O + O_2_ cultures than in the O_2_-replete CH_4_ + O_2_ cultures (Table 2), further demonstrating the beneficial contribution of N_2_O reduction for growth on CH_4_ at suboxic conditions.

### Transcriptomics

To study the expression of genes encoding respiration and C1-metabolism, we analyzed transcriptomes for strain T4 and IT6 grown on methanol under O_2_-replete (CH_3_OH + O_2_) and anoxic (CH_3_OH + N_2_O) conditions. In addition, we analyzed strain T4 grown on CH_4_ under O_2_-replete (CH_4_ + O_2_) and suboxic (CH_4_ + O_2_ + N_2_O) conditions. The overall regulation of key genes involved in denitrification and methane oxidation is depicted in Fig. 6 and Table 3, as well as in the supplementary material (Supplementary Figs. 8, 9, Supplementary Tables 5, 6). Differences in expression were considered significant if the Log_2_FC was higher than [0.85] or lower than [-1.0] with an adjusted p-value ≤ 0.05.

**Fig. 6.**
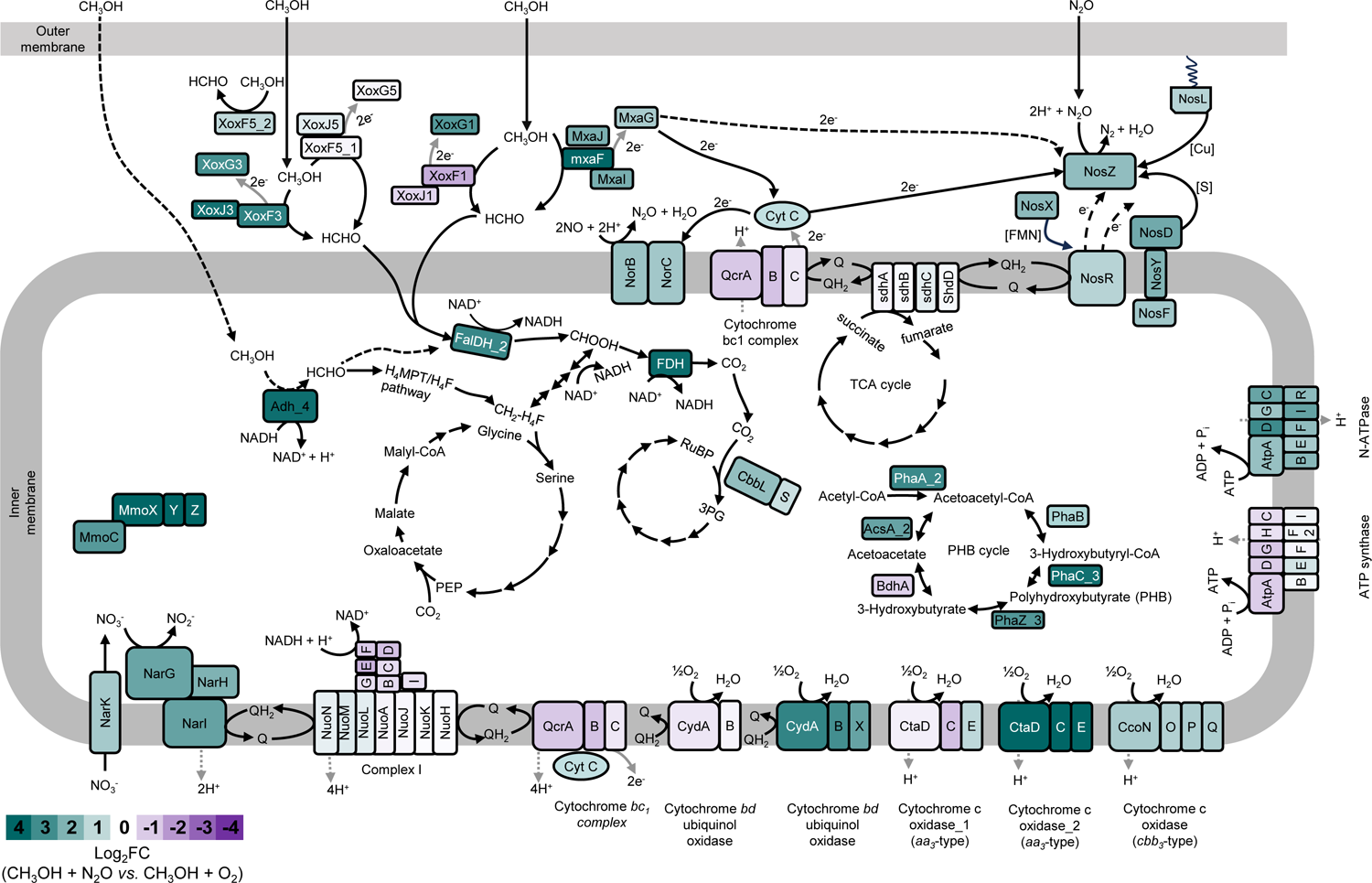
Metabolic reconstruction and transcriptional response of *Methylocella tundrae* T4 cells to O_2_-replete (CH_3_OH + O_2_) and anoxic (CH_3_OH + N_2_O) methanol-oxidizing growth conditions. The genes used to reconstruct the metabolic pathway are listed in Table S5. The gene products are shaded according to the relative fold change (Log_2_FC) in gene expression between cells grown under anoxic (CH_3_OH + N_2_O) and O_2_-replete (CH_3_OH + O_2_) conditions. Genes up-regulated in CH_3_OH + N_2_O-grown cells are shown in teal green, while those up-regulated in CH_3_OH + O_2_-grown cells are shown in purple. Note that proteins are not drawn to scale. **Methanol oxidation**: Methanol is oxidized to formaldehyde in the periplasmic space by the PQQ-dependent methanol dehydrogenase (Xox- and Mxa-type), T4_03519–21, T4_00353– 55, and T4_01862–76. The NAD(P)^+^-dependent alcohol dehydrogenase (T4_03199) may also be involved in methanol oxidation to formaldehyde in the cytoplasmic space during anaerobic growth on methanol. Formaldehyde oxidation to formate then proceeds via the tetrahydromethanopterin (H_4_MPT) pathway, and C1 incorporation into the serine cycle is mediated by the tetrahydrofolate (H_4_F) carbon assimilation pathway. The Calvin-Benson-Bassham pathway is also a possible route for CO_2_ fixation. **Nitrous oxide reduction**: N_2_O is reduced to N_2_ through the activity of nitrous oxide reductase in the periplasmic space. Electron transfer to NosZ occurs via cytochrome c from the cytochrome bc1 (Qcr) complex. Electron transfer to the NosZ may also involve direct interaction with methanol dehydrogenase C-type cytochrome (XoxG, MxaG). The NosR protein may be involved in the transfer of electrons to NosZ. The N_2_O reduction pathway is adapted from Hein and Simon ^139^ and Torres *et al*. ^99^.

**Table 3:**
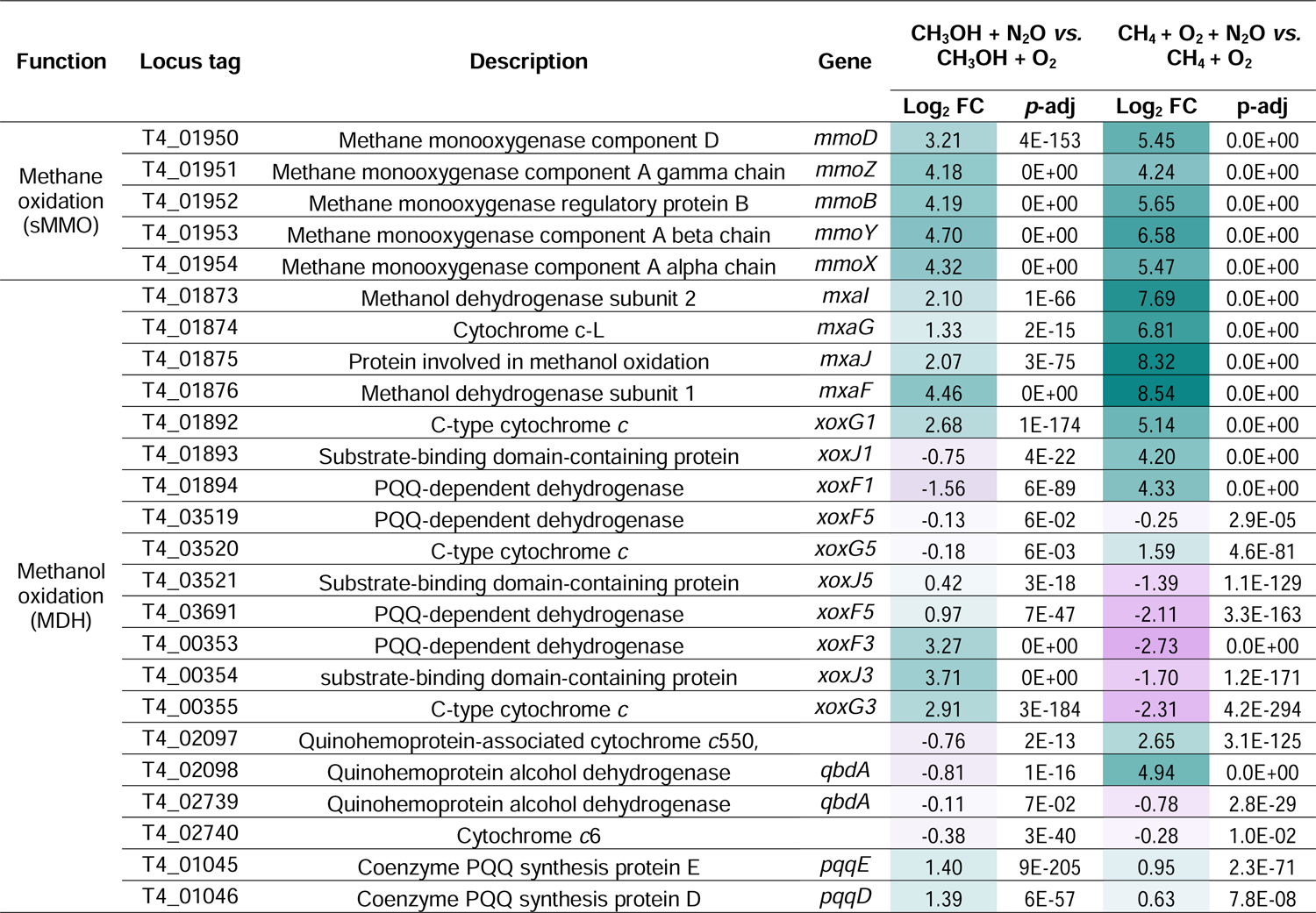
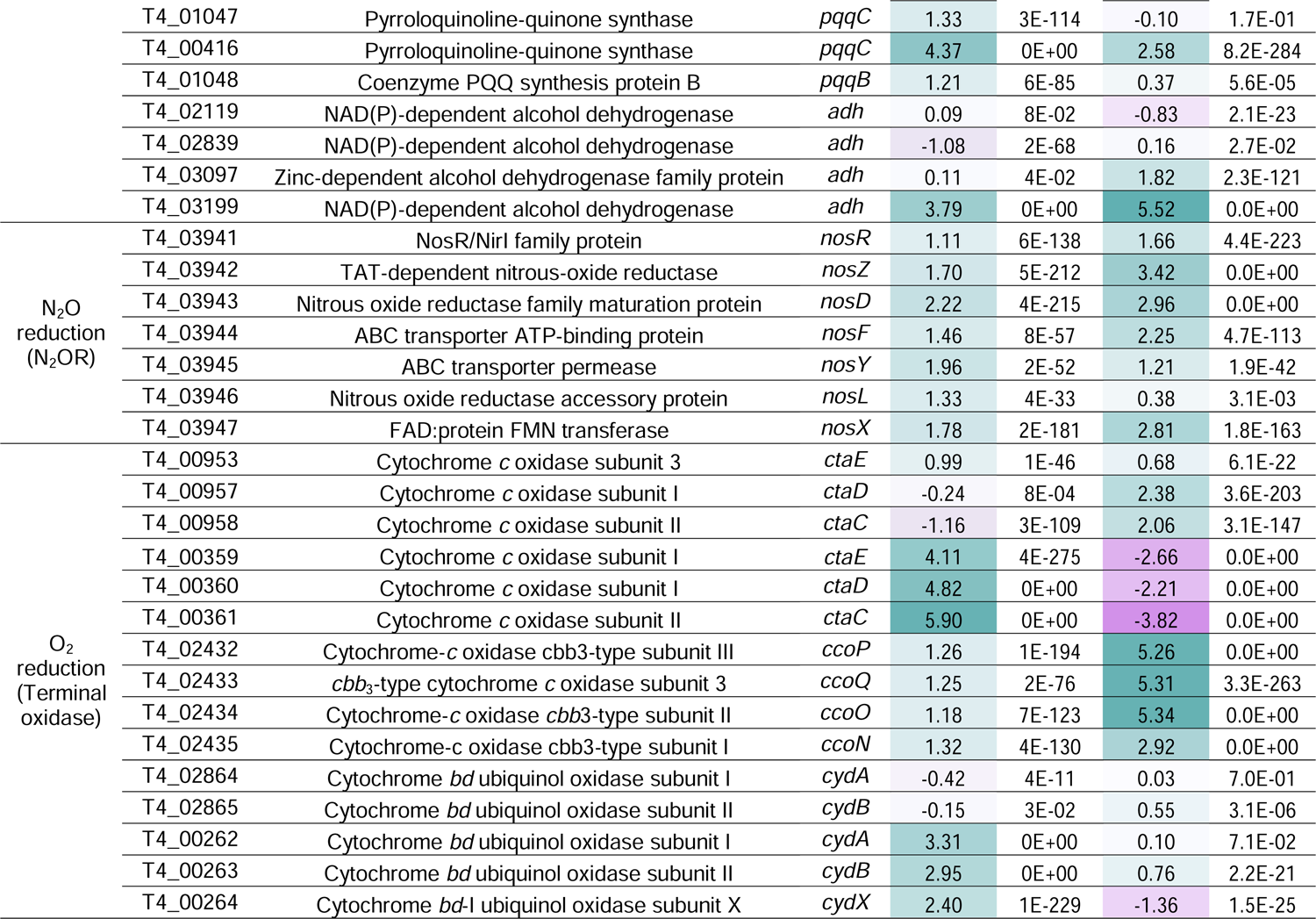
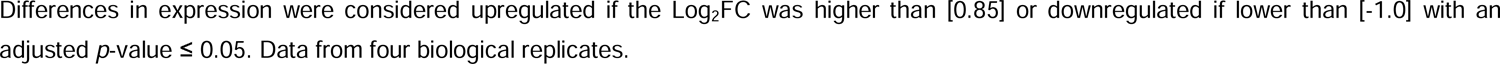
Transcriptional regulation of selected key genes in *Methylocella tundrae* T4 cells grown under methanol-(anoxic CH_3_OH + N_2_O versus O_2_-replete CH_3_OH + O_2_) and methane-oxidizing (suboxic CH_4_ + O_2_ + N_2_O versus O_2_-replete CH_4_ + O_2_) growth conditions.

#### N_2_OR (O_2_ replete vs. anoxic conditions)

The transcript levels of the N_2_OR-encoding genes (T4_03941– 47), *nosRZDFYLX*, were higher in strain T4 cells respiring N_2_O in the anoxic CH_3_OH + N_2_O conditions compared to strain T4 cells respiring O_2_ in the O_2_-replete CH_3_OH + O_2_ conditions (Supplementary Table 5). Under anoxic CH_3_OH + N_2_O conditions, expression of these genes increased 2- to 4.7-fold (Fig. 6, Table 3, Supplementary Table 5). Cells of strain IT6 respiring N_2_O in the anoxic CH_3_OH + N_2_O conditions showed transcriptional upregulation (1.9–6.7-fold) of four *nos* genes (*nosC1BZC2*) under anoxic conditions (Supplementary Fig. 9, Supplementary Table 6). Other *nos* operon genes (*nosYFDL*; IT6_00904–11) were expressed constitutively under both the anoxic CH_3_OH + N_2_O and O_2_-replete CH_3_OH + O_2_ conditions. The NosC1 and NosC2 proteins of *Wolinella succinogenes* were predicted to facilitate electron transfer from menaquinol to the periplasmic NosZ during the reduction of N_2_O to N_2_ ^61^ and are likely to play a similar role in strain IT6. Although the exact function of NosB has yet to be elucidated, Hein and colleagues ^96^ used a non-polar *nosB* deletion mutant of *Wolinella succinogenes* to show that it is necessary for N_2_O respiration. Overall, increased expression of N_2_OR-encoding genes in *Methylocella tundrae* T4 and *Methylacidiphilum caldifontis* IT6 cells during anaerobic growth indicates that the N_2_OR is functional in these methanotrophs and supports their ability to respire and grow using N_2_O as a terminal electron acceptor.

#### N_2_OR (O_2_ replete vs. suboxic conditions)

Transcript levels of N_2_OR-encoding genes were 2–10.7-fold higher in strain T4 cells grown under suboxic CH_4_ + O_2_ + N_2_O conditions than in cells grown under O_2_-replete CH_4_ + O_2_ conditions (Table 3, Supplementary Fig. 8, Supplementary Table 5). This finding is consistent with the N_2_O respiration activity and growth of strain T4 under suboxic CH_4_ + O_2_ + N_2_O conditions (Fig. 5), in which the cells can efficiently oxidize more CH_4_ (see Table 2), most likely leading to increased O_2_ use for CH_4_ oxygenation and the use of N_2_O for cellular respiration.

#### Other denitrification genes (O_2_ replete vs. anoxic/suboxic conditions)

Interestingly, even though N_2_O was the only electron acceptor present and NH_4_ ^+^ was the only nitrogen source provided, other denitrification enzyme genes were also upregulated (Fig 6, Supplementary Fig. 8, Supplementary Table 5) in *Methylocella tundrae* T4 cells grown on CH_4_ and CH_3_OH in response to N_2_O addition in suboxic and anoxic conditions, respectively. The expression of genes encoding respiratory NAR (*narGHJI*; 4–102-fold upregulation) and NOR (*norBC*; 2.6–58-fold upregulation) in strain T4 cells grown on CH_4_ and CH_3_OH was greatly increased in anoxic and suboxic N_2_O-respiring conditions. Similarly, in *Methylacidiphilum caldifontis* IT6, a gene NIR (*nirK*; IT6_01798) and genes encoding NOR (*norBC*; IT6_01320–1) were about 2-fold and 3.5-fold upregulated in the N_2_O-respiring anoxic CH_3_OH + N_2_O conditions, respectively, compared to the O_2_-replete CH_3_OH + O_2_ conditions (Supplementary Fig. 9, Supplementary Table 6) in the absence of NO_3_ ^−^ or NO_2_ ^−^ in the growth medium. Low oxygen and the presence of NO_3_ ^−^ and NO_2_ ^-^ as terminal electron acceptors are necessary for the expression of the genes of the denitrification pathway ^97–99^. Since neither NO_3_^−^ nor NO_2_^−^ were added to the cultures, the upregulation of NAR genes in strain T4 was probably influenced by the low O_2_ concentrations and/or presence of N_2_O. Although there is little evidence to support the upregulation of denitrification genes by N_2_O, in *Wolinella succinogenes*, an N_2_O reducer, the periplasmic nitrate reductase (Nap), cytochrome *c* nitrite reductase (Nrf), and N_2_O reductase (Nos) were upregulated in response to the presence of N_2_O ^100^. In this strain, N_2_O likely induced the Nap and Nrf systems independently even without notable amounts of NO_3_ ^−^ or NO ^100^. Significant upregulation of genes encoding NOR (Supplementary Table 5) and NOR regulatory proteins (Supplementary results) in strain T4 might be related to RNS detoxification (see above).

#### Methanol dehydrogenase (O_2_ replete vs. anoxic conditions)

In methanotrophs, methanol oxidation occurs in the periplasmic space by PQQ (pyrroloquinoline quinone)-dependent methanol dehydrogenase (MDH). Seven copies of the PQQ-dependent alcohol dehydrogenases (ADHs) ^101^ could perform this role in the genome of strain T4 (Supplementary Table 5). Five copies are type I ADHs (quinoproteins); one is a calcium-dependent MDH (MxaF-type MDH), and four are lanthanide-dependent MDHs (XoxF-type MDH), divided into clades 1 (XoxF1), 3 (XoxF3), and 5 (XoxF5; 2 copies) (Supplementary Fig. 10). The other two copies are type II ADHs (quinohemoproteins). In addition to the PQQ-dependent ADH, *Methylocella tundrae* T4 and *Methylacidiphilum caldifontis* IT6 genomes contained genes encoding cytosolic Zn^2+^-dependent ADH, which are part of a large family of enzymes that oxidize alcohols to aldehydes or ketones and reduce NAD(P)^+^ or a similar cofactor ^102, 103^ (Table 3, Supplementary Table 5).

Among the four XoxF-type MDHs encoded in the genome of strain T4, genes in a *xoxFGJ* operon (T4_03519–21), which included a gene encoding a XoxF5 enzyme, were found to be constitutively transcribed at high levels in cells grown under both O_2_-replete CH_3_OH + O_2_ and anoxic CH_3_OH + N_2_O conditions (Fig. 6, Table 3, Supplementary Table 5). Thus, the *xoxF5* gene likely encodes the predominant MDH used by strain T4 in both O_2_-respiring and N_2_O-respiring cells. The other singleton *xoxF*5 gene (T4_03691) and a *xoxF3* gene found in a separate *xoxFGJ* cluster (T4_00353–55) were also significantly upregulated in cells grown under anoxic CH_3_OH + N_2_O conditions in comparison to cells grown under O_2_-replete CH_3_OH + O_2_ conditions (Fig. 6, Table 3). Furthermore, we observed a significant upregulation (2- to 22-fold) of the genes encoding MxaFI-type MDH (T4_01872–76) in the anoxic CH_3_OH + N_2_O-grown cells (Fig. 6, Table 3). Thus, our results indicate the use of various MDHs by strain T4 during growth at anoxia. In strain IT6 a *xoxF* gene encoding a XoxF2-type MDH is present as part of the *xoxGJF* operon (IT6_00336–8) (Supplementary Table 6) and the expression of the *xoxF*2 gene was 2-fold upregulated in the N_2_O-respiring cells (Supplementary Fig. 9, Supplementary Table 6).

A cytosolic Zn^2+^-dependent ADH bound to NAD(P)^+^ is known to perform the oxidation of methanol only in Gram-positive methylotrophs ^104^. A Zn^2+^-dependent ADH (T4_03199) of strain T4 was significantly upregulated (13.8-fold) in the anoxic CH_3_OH + N_2_O-grown cells compared to the O_2_-replete CH_3_OH + O_2_-grown cells (Fig. 6, Table 3). The strain IT6 genome also contained three copies of genes encoding enzymes annotated as Zn^2+^-dependent ADH (Supplementary Table 6). The expression of two of these genes (IT6_01501 and IT6_01931) was found to be 3.9- and 2.5-fold, respectively, upregulated in N_2_O-respiring cells compared to cells respiring O_2_ (Supplementary Fig. 9, Supplementary Table 6). Even though PQQ-dependent MDHs have a high affinity for and activity with methanol as a substrate, their use in strictly anoxic conditions will be limited because PQQ biosynthesis requires molecular oxygen ^105, 106^. Thus, PQQ-dependent MDHs are suggested to be functional at completely anoxic conditions only when PQQ is carried over or provided externally ^107, 108^. On the other hand, Zn^2+^-dependent MDHs have the advantage of utilizing a ubiquitous cofactor, NAD(P)^+^, and can be functional during anaerobic growth ^109, 110^. This finding raises the possibility that strains T4 and IT6 can employ alternative ADHs such as the Zn^2+^-dependent ADH to facilitate methanol oxidation in strict anoxia. Some genes required for the following steps of methanol oxidation, formaldehyde, and formate dehydrogenases, were upregulated in strain T4, which are depicted in Fig. 6 and supplementary materials (Supplementary Fig. 8, Supplementary Table 5).

#### Methanol dehydrogenase (O_2_ replete vs. suboxic conditions)

Furthermore, we also examined expression levels of genes encoding MDHs in strain T4 cells grown under suboxic CH_4_ + O_2_ + N_2_O conditions (Table 3, Supplementary Fig. 8, Supplementary Table 5). Genes in the cluster T4_01862–76, which encodes the calcium-dependent MDH (MxaF-type MDH), had the highest transcript expression among all MDH-encoding genes in CH_4_-oxidizing cells grown under suboxic CH_4_ + O_2_ + N_2_O conditions. When compared to O_2_-replete CH_4_ + O_2_ conditions, the expression of genes within this cluster was 1.8- to 371.5-fold upregulated (Table 3, Supplementary Fig. 8, Supplementary Table 5). This is unexpected since typically, genes encoding the Mxa-type MDH are always downregulated in the presence of lanthanides ^111, 112^; which we also included (2 µM each of cerium and lanthanum) in the growth medium. Their apparent upregulation (even when lanthanides are present) suggests that this enzyme might play an important role in CH_4_ metabolism in the presence of N_2_O and suboxic conditions. As observed above, genes in the *xoxFGJ* operon (T4_03519–21) were also highly expressed at the suboxic conditions (Table 3, Supplementary Fig. 8, Supplementary Table 5), implicating a key MDH used by strain T4 in all three conditions. Genes in the cluster T4_01892–94 including the gene encoding the XoxF1 MDH were also significantly upregulated (18- to −35-fold) in the suboxic CH_4_ + O_2_ + N_2_O-grown cells compared to the O_2_-replete CH_4_ + O_2_-grown cells. The operon T4_02097–98, which encodes a cytochrome *c*550 (T4_02097) and a type II ADH (T4_02098), exhibited 6-fold and 30.6-fold upregulation, respectively, in cells grown under suboxic CH_4_ + O_2_ + N_2_O conditions as opposed to cells grown under O_2_-replete CH_4_ + O_2_ conditions. In addition, two Zn^2+^-dependent ADHs (T4_03097 and T4_03199) were significantly upregulated (3.5-fold and 46-fold, respectively) in strain T4 cells grown under suboxic CH_4_ + O_2_ + N_2_O conditions compared to cells grown under O_2_-replete CH_4_ + O_2_ conditions. Overall, it appears that cells oxidizing methanol under anoxia (CH_3_OH + N_2_O-grown cells) or those oxidizing methane under suboxia (CH_4_+O_2_+N_2_O-grown cells) use a distinct set of MDHs during N_2_O respiration than they do during O_2_ respiration.

#### Methane monooxygenase

The genomes of *Methylocella tundrae* T4 and *Methylacidiphilum caldifontis* IT6 contain genes that encode sMMO and pMMO, respectively. In the anoxic CH_3_OH + N_2_O conditions, genes encoding subunits of the soluble sMMO in T4 were 2–26-fold upregulated compared to O_2_-replete CH_3_OH + O_2_ conditions (Fig. 6, Table 3). In the suboxic CH_4_ + O_2_ + N_2_O conditions, all the genes (*mmoXYBZDCRG*) in the gene cluster T4_01946–54 displayed a high degree of transcriptional upregulation (18.7–96-fold) (Table 3, Supplementary Fig. 8, Supplementary Table 5). Similarly, in strain IT6, genes encoding subunits of the pMMO1 operon (*pmoCAB1*; IT6_01985–7) displayed the highest degree of transcriptional upregulation (∼94–749-fold) in the anoxically grown cells (CH_3_OH + N_2_O) (Supplementary Fig. 9, Supplementary Table 6).

The high upregulation of both monooxygenase-encoding operons under fully anoxic CH_3_OH + N_2_O conditions was surprising because CH_4_ was not used as an electron donor in anoxic conditions. Previous studies on verrucomicrobial methanotrophs have shown that the expression of genes encoding the *pmoCAB1* ^113, 114^ was upregulated at low O_2_ levels. In addition, *Methylosinus trichosporium* OB3b sMMO activity and protein expression were found to be significantly reduced under elevated O_2_ conditions (188 µM) compared with hypoxic conditions (24 µM) ^115^. Furthermore, elevated O_2_ levels in contrast to low O_2_ levels were found to impair the catalytic activity of *Methylosinus trichosporium* OB3b sMMO in the degradation of dichloroethane ^116^. Thus, in methanotrophs, upregulation of methane monooxygenase genes under O_2_ limiting conditions might be a strategy to produce more methane monooxygenase. This will lead to increased methane oxidation and thus provide stronger competition for the limited O_2_ with the terminal oxidase. Aside from the methane monooxygenase genes, group II and III truncated hemoglobin encoding genes were upregulated in *Methylocella tundra* T4 (T4_02445, T4_02637, and T4_00400; 4- to 12-fold) and *Methylacidiphilum caldifontis* IT6 (IT6_00149; 3-fold) cells in response to suboxia or anoxia (Supplementary Table 5, Supplementary Table 6). These truncated hemoglobins are thought to transport O_2_ to the methane monooxygenase ^27^.

#### Terminal oxidase

Three types of terminal oxidases (TOs) are encoded in the genome of strain T4, and their gene expression pattern is presented in Table 3 and Supplementary Table 5. Among these are one *cbb*_3_ (high-affinity TO), two *bd* (high-affinity TO), and two *aa*_3_ (low-affinity TO) ^117^. In the case of strain IT6, three types of TOs are encoded in its genome (Supplementary Table 6), as in most verrucomicrobial methanotrophs ^118^. They include one *cbb*_3_ (high-affinity TO), two *ba*_3_ (medium-affinity TO), and two *aa*_3_ (low-affinity TO) ^117, 119^. Due to the presence of multiple types of TOs, both strains T4 and T6 are well-suited to environments with fluctuating oxygen concentrations including peatlands, paddy soils, sulfidic oxygen-limited volcanic and geothermal ecosystems.

Strain T4 cells oxidizing methanol and respiring N_2_O in anoxic CH_3_OH + N_2_O conditions expressed all the genes encoding all terminal oxidases. Of these TOs, expression of genes encoding two of its high-affinity TOs, *bd*-type oxidase (T4_00262–66) and *cbb*_3_-type oxidase (T4_02432–35) were significantly upregulated 6–10-fold and ∼2.5-fold, respectively, in the anoxic CH_3_OH + N_2_O-grown cells in comparison to the O_2_-replete CH_3_OH + O_2_-grown cells (Fig. 6, Table 3). Interestingly, the anoxic CH_3_OH + N_2_O-grown cells also significantly upregulated the expression of genes encoding one of its low-affinity *aa*_3_-type oxidase (T4_00359–61) by 17- to 60-fold in comparison to the O_2_-replete CH_3_OH + O_2_-grown cells (Fig. 6, Table 3) as well. In the case of strain IT6 cells, transcriptional levels for genes encoding the high-affinity *cbb*_3_ TO were not significantly different between cells grown under O_2_-replete CH_3_OH + O_2_ and anoxic CH_3_OH + N_2_O conditions (Supplementary Fig. 9, Supplementary Table 6). There was a 6- to 30-fold upregulation of the expression of the genes encoding the three subunits of *ba*_3_ medium-affinity TO (IT6_00283-5) in anoxic CH_3_OH + N_2_O-grown cells (Supplementary Fig. 9, Supplementary Table 6). Also upregulated in the anoxic CH_3_OH + N_2_O-grown cells (2.8–172-fold) were genes for low-affinity *aa*_3_-type TO in the cluster IT6_00893–99, subunit I of *aa*_3_-type TO (*ctaD*; IT6_01499), and an accessory protein of *aa*_3_-type TO (*ctaA*; IT6_02050) (Supplementary Fig. 9, Supplementary Table 6).

Further, the transcript levels of the TO-encoding genes were compared in strain T4 cells grown under suboxic CH_4_ + O_2_ + N_2_O and O_2_-replete CH_4_ + O_2_ conditions (Supplementary Table 5). The genes in the operon T4_02429–35 encode a high-affinity *cbb*_3_-type oxidase that was upregulated 4 to 40-fold in suboxic CH_4_ + O_2_ + N_2_O-grown cells compared to the O_2_-replete CH_4_ + O_2_-grown cells (Table 3, Supplementary Fig. 8). In addition, the suboxic CH_4_ + O_2_ + N_2_O-grown cells increased the expression of some genes encoding a low-affinity *aa*_3_-type oxidase (T4_00956–58) by 2.6 to 5-fold (Table 3, Supplementary Fig. 8).

The upregulation of genes encoding high-affinity *cbb*_3_-type and *bd*-type TOs in suboxic and anoxic conditions by T4 strain cells, respectively, is consistent with studies indicating that these oxidases are used by microorganisms to conserve energy in low-oxygen environments ^117, 120, 121^. On the other hand, the observed expression and upregulation of genes encoding low-affinity TOs in strain T4 at suboxic condition is unexpected. However, some evidence suggests this is not uncommon: analysis of the transcriptome of *Methyloprofundus* sp. INp10, a methanotrophic endosymbiont of deep-sea bathymodiolin mussels, revealed the *aa*_3_ low affinity-TO as the most used TO in hypoxia ^122^. Some *Acidobacteria* strains were found to grow in suboxic conditions with low-affinity TOs at O_2_ concentrations as low as 1 nM ^123^. Likewise, the low-affinity *aa*_3_-type TO of the ammonia-oxidizing bacterium, *Nitrosomonas europaea*, was significantly upregulated during suboxic growth conditions ^124^. Oxygen deprivation led to an increase in the expression of genes encoding low-affinity TOs in both the T4 and IT6 strains. Because low-affinity TOs are more efficient at generating ATP than high-affinity TOs ^69, 125, 126^, their utilization may be ecologically beneficial to microorganisms in low-oxygen ecosystems. Furthermore, the upregulation of low-affinity TOs coincided with the upregulation of denitrification genes, including the N_2_OR genes, which also coincided with the upregulation of methane monooxygenase. Thus, especially in methanotrophs, the upregulation of low-affinity TOs in suboxic conditions may help the redistribution of the limited O_2_ to the methane monooxygenase especially in the presence of other terminal electron acceptors, such as N_2_O.

## Conclusions

In this study, we revealed that sMMO- and pMMO-containing acidophilic methanotrophs of the genera *Methylocella* and *Methylacidiphilum* can grow anoxically by respiring N_2_O using clade I and II NosZ, respectively. N_2_O reduction can be expanded to an extremely acidic pH of 2.0. Further, a higher growth yield during methanotrophic growth at O_2_-limiting conditions by utilization of N_2_O can provide a competitive advantage. This study significantly expands our perception of the potential ecological niches of aerobic methanotrophs. In addition to mitigating CH_4_ and CO_2_ emissions, aerobic methanotrophs potentially play a role in reducing the emission of the climate-active and ozone-depleting gas, N_2_O, providing application potential in controlling the discharge of these key GHGs from natural and engineered systems.

## Materials and Methods

### Bacterial strains and growth conditions

The methanotrophic bacterial strains used for the experiments include *Methylacidiphilum caldifontis* IT6, *Methylacidiphilum infernorum* IT5, *Methylocella tundrae* T4 (= KCTC 52858^T^), *Methylocella silvestris* BL2 (= KCTC 52857^T^), *Methylocystis* sp. SC2 ^127^ (kindly provided by Dr. Angela Smirnova and Prof. Peter F. Dunfield, Department of Biological Sciences, University of Calgary, Canada), and three in-house *Methylocystis echinoides*-like isolates (strains IM2, IM3, and IM4). The *Methylacidiphilum* strains are also in-house strains isolated previously from a mud-water mixture taken from Pisciarelli hot spring in Pozzuoli, Italy ^56^. The *Methylocella* strains were obtained from the Korean Collection for Type Cultures (KCTC). Growth of the bacterial strains was performed using a low salt mineral (LSM) medium described previously ^56^. The pH of the medium was adjusted to pH 2.0 with concentrated sulfuric acid (filter-sterilized) for the *Methylacidiphilum* strains and to pH 5.5 with 20 mM 2-morpholinoethanesulfonic acid (filter-sterilized) for the *Methylocella* strains. The cultures were incubated at 52 °C for *Methylacidiphilum* strains (IT5 and IT6) and 28 °C for *Methylocella* strains (T4 and BL2) with shaking at 160 rpm. Unless stated otherwise, ammonium sulfate, (NH_4_)_2_SO_4_, was used as the nitrogen source.

### Genomic and phylogenetic analysis

To analyze the distribution of denitrification genes in methanotrophs, isolate or metagenome-assembled genomes (MAG) (meeting the following CheckM criteria: completeness > 60% and contamination < 10%) were obtained from the NCBI assembly database. The extraction of high-molecular-weight genomic DNA from *Methylocella tundrae* T4 and the in-house *Methylocystis* isolates (strains IM2, IM3, and IM4), as well as their genome sequencing and annotation, are described in the *Supplementary Information*, *DNA Isolation, and Genomic Analysis*. Genomes were manually annotated with the Prokka annotation pipeline (v1.14.6) with the default option ^128^. The annotated protein sequences were re-annotated against the NCyc ^129^ and BV-BRC ^130^ databases. Based on the annotation results, genes encoding enzymes of the denitrification pathway (*napA*, *napB*, *narG*, *narH*, *narI*, *nirS*, *nirK*, *norB*, *norC*, and *nosZ*) were selected. The obtained protein sequences were aligned with reference sequences downloaded from the NCBI protein and nucleotide databases, and their identity was assured using tree-building procedures; only candidate genes that clustered with reference sequences were considered true hits and counted.

For phylogenetic analyses of the NosZ proteins and methanol dehydrogenases of strains T4 and IT6, representative amino acid sequences of the genes of related taxa were obtained from NCBI. The derived amino acid sequences of the NosZ and methanol dehydrogenases (XoxF and MxaF) were aligned using MAFFT ^131^. Maximum-likelihood trees were inferred with IQ-TREE. The constructed trees and operon arrangements were visualized using iTOL and used for annotation. Genomic islands were predicted using the IslandViewer web server ^132^.

### Anoxic growth coupled with N_2_O reduction

To demonstrate the ability of N_2_OR-containing methanotrophs to grow using N_2_O as the electron acceptor, we established anoxic batch cultures of *Methylocella tundrae* T4 and *Methylacidiphilum caldifontis* IT6 in 160-mL bottles containing 20 mL of LSM media and inoculated with 1–5% (v/v) actively growing-cells from the log phase (starting OD_600_ values ≤ 0.05). To remove oxygen, nitrogen gas (N_2_, purity >99.999%) was introduced into the bottles via a long needle (18G). Following that, the bottles were flushed with N_2_ gas for 20 minutes before being sealed with gas-tight butyl rubber stoppers and aluminum crimp seals to prevent O_2_ leakage. We used contactless trace-range oxygen sensor spots (TROXSP5) to monitor O_2_ contamination (< 0.10%, v/v) in the culture bottles incubated after N_2_-flushing (see *Supplementary Information*, *Analytical methods*, for details). These spots have a detection limit of 20 nM O_2_. Chemical-reducing agents, Na_2_S (0.5, 1, and 2 mM), cysteine (0.5 mM), DTT (0.5 mM), and titanium citrate (0.5 and 1 mM) in the media resulted in severe cell toxicity, hindering their use in this study as previously reported for N_2_OR reducer Anaeromyxobacter dehalogenans ^133^. When the cultures were incubated without the chemical-reducing agents, the cells completely depleted the trace O_2_ concentration present in the culture bottles in less than 24 hours as measured by the oxygen sensor spots.

N_2_OR-lacking close relatives, *Methylocella silvestris* BL2 and *Methylacidiphilum infernorum* IT5, which are closely related to *Methylocella tundrae* T4 and *Methylacidiphilum caldifontis* IT6, respectively, were used as negative controls. Methanol (30 mM), N_2_O (5%, v/v), and CO_2_ (5%, v/v) were used as the energy source, electron acceptor, and carbon source, respectively. In addition, pyruvate (10 mM) and hydroxyacetone (acetol) (10 mM) were tested as the sole C-C electron donors in strains T4 and IT6, respectively. Furthermore, strain IT6 cells were investigated to grow chemolithoautotrophically in sealed 1-liter bottles (duplicate) containing 20 mL of LSM medium at pH 2.0 on H_2_ (10% v/v) with and without N_2_O (5% v/v). As part of the control experiments, we incubated cells from the four strains in LSM media under anoxic conditions (without N_2_O) to assess the contribution of the initial trace O_2_ present in the culture bottles to biomass increase. The increase in biomass as OD_600_ by the trace O_2_ in the control cultures was negligible when compared to cultures growing with N_2_O as the sole electron acceptor (see Fig. 2c, 2f, 2i, 2l). Positive control experiments with methanol (30 mM) and O_2_ (5%, v/v) as the electron donor and electron acceptor, respectively, were conducted for each strain. The concentrations of H_2_, O_2_, N_2_O, NO_3_ ^−^, and NO_2_ ^−^ concentrations were monitored at intervals during incubations (described in the *Supplementary Information*, *Analytical methods*). Cell growth was also evaluated using optical density measurements (λ=600 nm). All growth experiments were performed in triplicates unless otherwise stated.

Next, we checked the anoxic growth of *Methylocella* strains on NO_3_ ^−^ (2 to 4 mM KNO) as the terminal electron acceptor instead of N_2_O. Methanol (30 mM) was used as the sole electron donor and 2 mM NH_4_ ^+^ was used as the N-source. To compare the effect of electron donors on NO_3_ ^−^ and NO_2_ ^−^ reduction, *Methylocella* strains were also anoxically grown in LSM medium containing a C-C substrate, pyruvate (10 mM). Cells of strain T4 were grown under O_2_-replete (O_2_; 21%, v/v) or anoxic conditions (O_2_; 0%, v/v, N_2_O; 5%, v/v) for the NO_2_ ^−^ toxicity test (triplicates) with varying NO_2_ ^−^ (KNO_2_) concentrations (0, 0.01, 0.03, 0.1, 0.3, and 1 mM).

### Kinetic analysis using microrespirometry (MR)

For kinetic analysis using microrespirometry (MR), *Methylocella tundrae* T4 cells were grown under three different O_2_ conditions: O_2_-replete (CH_3_OH + O_2_), suboxic (CH_4_ + O_2_ + N_2_O), and anoxic (CH_3_OH + N_2_O). *Methylacidiphilum caldifontis* IT6 cells were grown under O_2_-replete (CH_3_OH + O_2_) and anoxic (CH_3_OH + N_2_O) conditions. The O_2_-replete growth conditions included ambient air (21% O_2_, v/v) and CH_3_OH (30 mM) as the sole electron donor. The suboxic cell cultures were grown under a condition that included CH_4_ (5%, v/v) as the sole electron donor and O_2_ (0.5%, v/v) with N_2_O (1%, v/v) as terminal electron acceptors. O_2_ (0.5%, v/v) was resupplied on an intermittent basis before its depletion. Anoxically grown cells were cultured in bottles containing 30 mM CH_3_OH as the sole electron donor and 5% (v/v) N_2_O as the terminal electron acceptor. The cultures were monitored daily and harvested as soon as active consumption of electron donors and acceptors was detected. After being collected by centrifugation (5000 × g, 30 min, 25 °C), the cells were washed twice with substrate- and N-source-free MES-buffered LSM (20 mM MES; pH 5.5) or H_2_SO_4_-buffered LSM (4 mM H_2_SO4; pH 2.0) and then resuspended in 20 mL of the same media without electron donors and acceptors. In the case of the anoxic and suboxic grown cells, the cell suspensions were transferred to sealed 20-mL bottles and flushed with nitrogen gas (N_2_, purity >99.999%) before use. The cell suspensions were dispensed into a double-port MR chamber (no headspace) with a capacity of 5 or 10 mL outfitted with O_2_ and N_2_O-detecting microsensors, two MR injection lids, and two glass-coated stir bars. Kinetics and stoichiometry of N_2_O and O_2_ reduction coupled to CH_3_OH oxidation were estimated using anoxic CH_3_OH + N_2_O- and oxic CH_3_OH + O_2_-grown cells, respectively. Anoxic CH_3_OH + N_2_O-grown cells were used to test CH_3_OH-dependent O_2_ and N_2_O uptake by strains IT6 (starting OD_600_ = 0.96) and T4 (starting OD_600_ = 0.79). The effect of O_2_ to N_2_OR activities of strains T4 and IT6 was determined by spiking varying O_2_ to the N_2_O respiring cells. In a 5-mL MR chamber, suboxic CH_4_ + O_2_ + N_2_O-grown cells of strain T4 (starting OD_600_ = 1.0) were used to test the CH_4_-dependent simultaneous respiration of O_2_ and N_2_O.

All MR experiments were performed in a recirculating water bath at 27 °C and 50 °C for strains T4 and IT6, respectively. A 10-µL or 50-µL syringe (Hamilton, Reno, USA) fitted with a 26G needle was used to inject the substrate (CH_4_, CH_3_OH, N_2_O, or O_2_) into the chamber via an injection port. Concentrations of O_2_ and N_2_O were measured using an OX-MR oxygen microsensor (OX-MR-202142, Unisense, Aarhus, Denmark) and a N_2_O-MR sensor (N2O-MR-303088, Unisense), respectively. The detection limits of the OX-MR and N_2_O-MR microsensors are 0.3 µM O_2_ and 0.1 µM N_2_O, respectively. The OX-MR and N_2_O-MR microsensors were directly plugged into a microsensor multimeter before being polarized for more than a day and calibrated according to the manufacturer’s instructions. All data from the microsensor multimeter was logged onto a laptop using SensorTrace Logger software (Unisense). Anoxically prepared aliquots of N_2_O, CH_4_, and CH_3_OH were injected into the MR chamber via the injection port with a 10-µL syringe (Hamilton, Reno, USA). Anoxic substrate-free LSM media (at pH 2.0 and 5.5) were prepared by sparging the solutions with N_2_ gas for 1 hour before use. Anoxic saturated-aqueous CH_4_ and N_2_O solutions were made in capped 160-mL bottles containing 100 mL of LSM medium and pressurized with CH_4_ or N_2_O (1, 2, or 3 atm; 100%, v/v). Saturated-aqueous O_2_ solutions were prepared in capped 160-mL bottles containing 100 mL of LSM medium and pressurized with O_2_ (1, 2, and 3 atm; 100%, v/v).

### Growth based on CH_4_ oxidation coupled with co-respiration of O_2_ and N_2_O

Suboxic cultivations were carried out to investigate the growth of *Methylocella tundrae* T4 by oxidizing methane with simultaneous respiration of O_2_ and N_2_O. The experiments were conducted in N_2_-flushed 2-liter sealed bottles (2,360 mL) containing 60 mL of LSM medium with 2 mM NH_4_ ^+^ as the N-source. The headspace of the bottles was composed of CH_4_ (5%, v/v), O_2_ (0.5%, v/v), N_2_O (1%, v/v), and CO_2_ (5%, v/v) and supplemented with additional O_2_ (∼0.5%, v/v) before its depletion. The headspace gas (CH_4_, N_2_O, and O_2_) mixing ratios were monitored at intervals during incubations as described in *Supplementary Information*, *Analytical Methods*. To investigate the growth benefits of cells of strain T4 respiring N_2_O in tandem with O_2_ during CH_4_ oxidation, an O_2_-replete culture was included for comparison (triplicates). The apparent increase in cell densities of both growth conditions was compared using optical density measurements (λ =600 nm).

### Transcriptome analysis

Cells of strains T4 and IT6 were cultured in 60 mL of LSM medium at pH 5.5 and pH 2.0 in sealed 2-liter bottles (4 or 5 replicates) for transcriptome analyses. Strain T4 cells were cultured under three different O_2_ levels, with the first setting being O_2_-replete (CH_4_ + O_2_ and CH_3_OH + O_2_), the second being suboxic (CH_4_ + O_2_ + N_2_O), and the third being anoxic (CH_3_OH + N_2_O). Strain IT6 was cultivated in O_2_-replete CH_3_OH + O_2_ and anoxic CH_3_OH + N_2_O conditions. Cells were grown anoxically in bottles containing 30 mM CH_3_OH as the sole electron donor and 5% N_2_O as the terminal electron acceptor. The O_2_-replete growth conditions were made up of ambient air (21% O_2_, v/v) with CH_4_ (5%, v/v) or CH_3_OH (30 mM) serving as the sole electron donor. The suboxic growth conditions were made up of a mixture of CH_4_ (5% v/v) as the sole electron donor and O_2_ (0.5% v/v) and N_2_O (1% v/v) as terminal electron acceptors. Before the depletion of O_2_, additional O_2_ was resupplied intermittently at a mixing ratio of 0.5% (v/v). Contactless trace-range oxygen sensor spots (TROXSP5) were installed into the culture bottles to monitor O_2_ concentration.

The cells were harvested during the mid-exponential phase at 5,000 × g for 10 minutes at 25 °C. Total RNA was extracted from the cells in four replicates using the AllPrep DNA/RNA Mini Kit (Qiagen) according to the manufacturer’s protocol. RNA quality was checked with the Agilent 2100 Expert Bioanalyzer (Agilent), and cDNA libraries were prepared from the RNA samples using the Nugen Universal Prokaryotic RNA-Seq Library Preparation Kit. The cDNA libraries were sequenced using NovaSeq6000 (Illumina) at LabGenomics (Seongnam, Korea). Read quality was evaluated with FastQC (v0.11.8) ^134^. Trimmomatic (v0.36) ^135^ was used to trim reads with the options: SLIDINGWINDOW:4:15 LEADING:3 TRAILING:3 MINLEN:38 HEADCROP:13. Reads mapped to strains T4 and IT6 rRNA sequences were removed with SortMeRNA (v2.1) ^136^. The remaining reads were aligned to the genomes of strains T4 and IT6 using Bowtie2 (v2.4.4) ^137^, and the reads mapped to each gene were counted using HTSeq (v0.12.3) ^138^. Expression values are presented as transcripts per kilobase million (TPM). The statistical analysis of differentially expressed genes was performed using the DESeq2 package in R.

## Data Availability

The complete genome sequence of strain T4 was deposited in the National Center for Biotechnology Information (NCBI) GenBank (accession nos. CP139087-CP139089). The whole transcriptome data were deposited in the NCBI BioProject database under the accession number PRJNA1050235.

## Acknowledgments

This work was supported by the NRF (National Research Foundation of Korea) grant funded by the Korean government (Ministry of Science and ICT) (2021R1A2C3004015) and Basic Science Research Program through NRF funded by the Ministry of Education (2020R1A6A1A06046235). J-HG was supported by the NRF grant funded by the Korean government (Ministry of Science and ICT) (RS-2023-00213601). M-YJ was supported by the NRF grant funded by the Korean government (Ministry of Science and ICT) (2021R1C1C1008303 and 2022R1A4A503144711)

## Author contributions

SIA, J-HG, and S-KR designed research. SIA, J-HG, M-YJ, and YK performed research. SIA, J-HG, YK, M-YJ, PFD, and S-KR analyzed data. SIA, J-HG, M-YJ, PFD, MW, and S-KR wrote the manuscript with contributions and comments from all co-authors.

## Competing interests

The authors declare no competing interests.

## Notes

### Competing Interest Statement

The authors have declared no competing interest.

