## Supplemental Information for "Nitrous oxide respiration in acidophilic methanotrophs"

*Sung-Keun Rhee

**The file includes:**

**Supplementary Figures and Tables**

Supplementary Materials and Methods
Supplementary Results
Figures S1 to S10
Legends for Tables S1 to S6
SI References

### Supplementary Materials and Methods

#### *Enrichment and isolation of Methylocystis strains*

The $N_2OR$-containing *Methylocystis* strains were isolated from an acidic forest soil in Chungcheongbuk-do, South Korea (36°55'31" N 127°54'86" E). The soil sample preparation and initial enrichment of the methanotrophs have previously been described [1]. Methanotrophic isolates were obtained by diluting the enrichment cultures repeatedly, as previously described (ref). Briefly, the most diluted culture exhibiting methane oxidation was serially diluted and filtered through 0.2-μm Track-Etch membrane polycarbonate filters (Whatman). The filters were placed on LSM medium (pH 5.5) in Petri dishes and incubated at 30°C in airtight containers containing $CH_4$ (10%, v/v) and $CO_2$ (5%, v/v). Colonies that appeared on the filters after 3 weeks of incubation were transferred to fresh LSM medium in 160-mL serum vials with the same gas composition. Three individual methanotrophic isolates were identified by sequencing the 16S rRNA gene with the 27F/1492R primer set [2]. The purity of the isolates was confirmed by seeding aliquots of the $CH_4$-grown cultures into the LSM medium with 0.05% (w/v) yeast extract, tryptic soy broth, and Luria-Bertani broth without $CH_4$ and incubating at 30 °C. Three methanotrophic isolates, IM2, IM3, and IM4, shared 99.46% 16S ribosomal RNA (rRNA) gene-sequence identity with the alphaproteobacterial methanotroph *Methylocystis echinoides* LMG27198. The three strains share average nucleotide identity values ranging from 81.85–81.93 with *Methylocystis echinoides* LMG27198, implying that they represent a new species in the genus *Methylocystis*.

#### *Analytical methods*

A YL 6100 gas chromatograph (YL Instrument Co., Anyang, South Korea) with a flame ionization detector (FID) and a thermal conductivity detector (TCD) was used to analyze the mixing ratios of $CH_4$, $N_2O$, and $H_2$ in the headspace of the sealed bottles used to cultivate the *Methylocella* and *Methylacidiphilum* strains. Using a Hamilton glass syringe, 100 µL of the sealed bottle headspaces were injected into a gas chromatograph equipped with MolSieve 5A column (3Ft, 1/8, 2mm, 60/80 SST, Agilent Technologies, Inc., CA, USA; for separating $H_2$, $O_2$, and $N_2O$) and Haysep N column (7Ft, 1/8, 2mm, 60/80 SST, Agilent Technologies, Inc., CA, USA; for separating $CO_2$ and $CH_4$) to determine the gases present. Helium was used as the carrier gas, with a flow rate of 15 $mL \cdot min^{-1}$. By utilizing pure gases of known concentrations, a calibration curve of the gases used as substrates was generated. The bottles were fitted with contactless trace range oxygen sensor spots (TROXSP5, PyroScience, Germany) calibrated at 0% and ambient air (21% oxygen), and a FireSting-Pro multi-analyte meter (FSPRO-4, PyroScience, Germany) was used to measure the $O_2$ concentration in the sealed bottles. Acidic Griess reagent and $VCl_2$/Griess reagent were used for photometric quantification of $NO_2^-$ and $NO_3^-$ concentrations [3], respectively, using a SpectraMax M2 microplate reader (Molecular Devices, USA).

#### *DNA isolation and genomic analysis*

High-molecular-weight genomic DNA was extracted from cultures of *Methylocella tundrae* T4 grown in methanol for 200 mL and the *Methylocystis* isolates (strains IM2, IM3, and IM4) grown in $CH_4$ using a modified CTAB method [4]. The genomes of *Methylocella tundrae* T4 and *Methylocystis* sp. IM3 were sequenced at LabGenomics (Seongnam, Republic of Korea) and Macrogen (Seoul, Republic of Korea) using the PacBio RS II (long-read sequencing) and Illumina HiSeq (2 x 150 bp) platforms, respectively. The genomes of *Methylocystis* sp. IM2 and *Methylocystis* sp. IM4 were sequenced using MinION R10.4.1 flow cell (Oxford Nanopore Technologies). The PacBio reads were assembled with the Trycycler pipeline (v0.5.4) [5]. Filtered reads were subsampled and assembled using Miniasm/Minpolish (v0.3-r179) [6], Flye (v2.9.3) [7], and Raven (v1.8.3) [8] assemblers. The consensus contigs were polished with Illumina short reads using Polypolish (v0.5.0) [9] and POLCA (v4.0.5) [10]. The circularity was confirmed during the Trycycler pipeline assembly and again by mapping the Illumina reads backward. *De novo* genome assembly of the MinION long reads was accomplished using the Canu-SMARTdenovo method [11]. Canu (v. 1.1.1) [12] was used to correct the reads before they were assembled with SMARTdenovo [13]. The accuracy of the assembled genome was improved by polishing with Nanopolish (v0.10.1) [14]. Annotation of methanotrophs' genomes was performed with the Prokka annotation pipeline (v1.14.6) [15], NCBI Prokaryotic Genome Annotation Pipeline (PGAP) [16], MicroScope [17], and PATRIC [18] annotation platforms. Functional assignment of the predicted genes was improved using a set of public databases (InterPro [19], GO [20, 21], PFAM [22], CDD [23], TIGRFAM [24], and EggNOG [25]). Prediction of signal peptides and transmembrane helices was performed using the web-based services SignalP (v5.0) [26] and TMHMM (v2.0) [27] with default settings.

### Supplementary Results

#### *$N_2O$ reduction kinetics*

$N_2O$ depletion in strains T4 and IT6 followed Michaelis-Menten kinetics (Fig. S5). We could obtain a maximum $N_2O$ reduction rate ($V_{max(app)}$) of 1.122 ± 0.005 mmol $N_2O \cdot h^{-1} \cdot g$ $DW^{-1}$ and an apparent $N_2O$ affinity ($K_{m(app)}$) of 5.937 ± 0.005 µM for the clade II $N_2O$-reducer, strain T4. The nitrous oxide reductase kinetics of strain IT6 cells showed a lower $N_2O$ reduction rate ($V_{max(app)}$ = 0.414 ± 0.003 mmol $N_2O \cdot h^{-1} \cdot g$ $DW^{-1}$) but a higher affinity constant value ($K_{m(app)}$) = 1.128 ± 0.043 µM $N_2O$). The affinity constant values of strain T4 and strain IT6 were consistent with those of clade I and clade II $N_2O$-reducers [28, 29], respectively.

#### *Expression of denitrification transcriptional regulators*

Regulatory proteins like the CRP/FNR family of transcriptional regulators, including FnrP (fumarate and nitrate reduction protein) and NNR (nitrite reductase and nitric oxide reductase regulator), respond to environmental signals like $O_2$ and NO to regulate the transcription and expression of denitrification genes [30, 31]. FnrP contains an oxygen-sensitive [4Fe-4S] cluster and regulates the oxygen-dependent transcriptional activation of many genes, including the *nar* operon [30]. Upstream of the *nos* genes of strain T4, we found three genes whose expression was upregulated (~ 2–29-fold) in suboxic $CH_4$ + $O_2$ + $N_2O$ and anoxic $CH_3OH$ + $N_2O$ conditions (Supplementary Table 5). These include T4_03938, which encodes the NNR; T4_03939, which encodes the FnrP; and T4_03940, which encodes the flavodiiron protein rubrerythrin involved in the reduction of $O_2$ and/or NO to $H_2O$ and $N_2O$ [32]. Two genes encoding the FnrP (T4_02964 and T4_02964), that activate NAR genes expression in response to $O_2$ deprivation [33] were found immediately downstream of the NAR genes, and their expressions were upregulated (~ 2–27-fold) in response to $N_2O$ respiration in suboxic and anoxic conditions (Supplementary Table 5). The expression of genes encoding NOR regulatory proteins (T4_00479–80), FnrP, and NNR, which are located immediately downstream of the *nor* genes, was also found to be significantly upregulated (14- to 21-fold) in the suboxic $CH_4$ + $O_2$ + $N_2O$ growth conditions (Supplementary Table 5). This suggests that the FnrP- and NNR-encoding genes are involved in upregulations of strain T4 denitrification genes in response to $O_2$ limitation as observed in *Paracoccus denitrificans* [33]. Overall, data from the transcriptome analysis reveals that the expression of denitrification genes in *Methylocella tundrae* T4 is regulated during $N_2O$ respiration in suboxic or anoxic conditions. The ability of strain T4 to upregulate the expression of denitrification genes in suboxia correlates well with its ability to employ a hybrid respiration system (i.e., simultaneous respiration of $O_2$ and $N_2O$) during $CH_4$ oxidation.

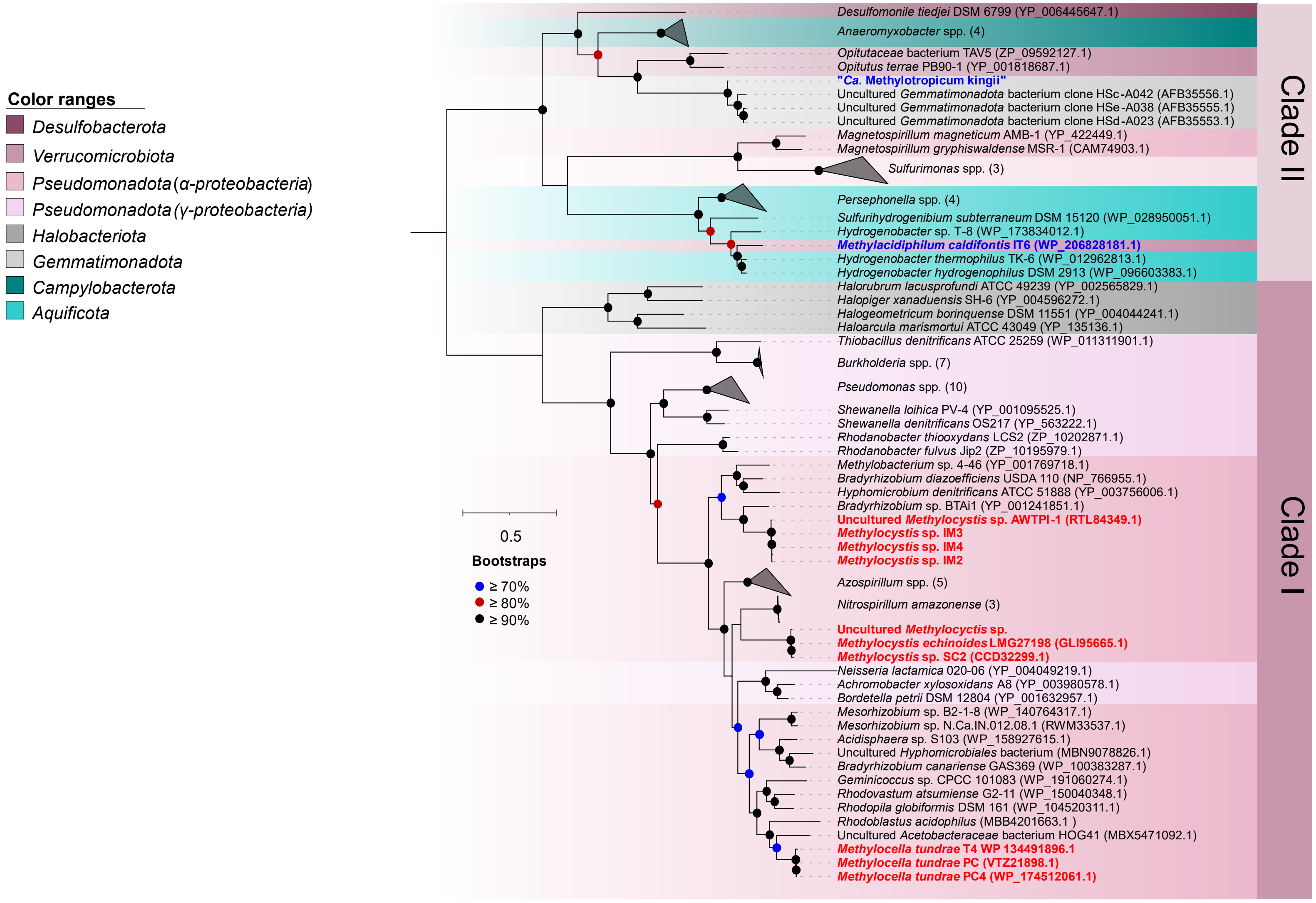

Color ranges
Desulfobacterota
Verrucomicrobiota
Pseudomonadota (α-proteobacteria)
Pseudomonadota (γ-proteobacteria)
Halobacteriota
Gemmatimonadota
Campylobacterota
Aquificota
0.5
Bootstraps
≥ 70%
≥ 80%
≥ 90%
Clade II
Clade I
Desulfomonile tiedjei DSM 6799 (YP_006445647.1)
Anaeromyxobacter spp. (4)
Opitutaceae bacterium TAV5 (ZP_09592127.1)
Opitutus terrae PB90-1 (YP_001818687.1)
"Ca. Methylotropicum kingii"
Uncultured Gemmatimonadota bacterium clone HSc-A042 (AFB35556.1)
Uncultured Gemmatimonadota bacterium clone HSe-A038 (AFB35555.1)
Uncultured Gemmatimonadota bacterium clone HSd-A023 (AFB35553.1)
Magnetospirillum magneticum AMB-1 (YP_422449.1)
Magnetospirillum gryphiswaldense MSR-1 (CAM74903.1)
Sulfurimonas spp. (3)
Persephonella spp. (4)
Sulfurihydrogenibium subterraneum DSM 15120 (WP_028950051.1)
Hydrogenobacter sp. T-8 (WP_173834012.1)
Methylacidiphilum caldifontis IT6 (WP_206828181.1)
Hydrogenobacter thermophilus TK-6 (WP_012962813.1)
Hydrogenobacter hydrogenophilus DSM 2913 (WP_096603383.1)
Halorubrum lacusprofundi ATCC 49239 (YP_002565829.1)
Halopiger xanaduensis SH-6 (YP_004596272.1)
Halogeometricum borinquense DSM 11551 (YP_004044241.1)
Haloarcula marismortui ATCC 43049 (YP_135136.1)
Thiobacillus denitrificans ATCC 25259 (WP_011311901.1)
Burkholderia spp. (7)
Pseudomonas spp. (10)
Shewanella loihica PV-4 (YP_001095525.1)
Shewanella denitrificans OS217 (YP_563222.1)
Rhodanobacter thiooxydans LCS2 (ZP_10202871.1)
Rhodanobacter fulvus Jip2 (ZP_10195979.1)
Methylobacterium sp. 4-46 (YP_001769718.1)
Bradyrhizobium diazoefficiens USDA 110 (NP_766955.1)
Hyphomicrobium denitrificans ATCC 51888 (YP_003756006.1)
Bradyrhizobium sp. BTAi1 (YP_001241851.1)
Uncultured Methylocystis sp. AWTPI-1 (RTL84349.1)
Methylocystis sp. IM3
Methylocystis sp. IM4
Methylocystis sp. IM2
Azospirillum spp. (5)
Nitrospirillum amazonense (3)
Uncultured Methylocyctis sp.
Methylocystis echinoides LMG27198 (GLI95665.1)
Methylocystis sp. SC2 (CCD32299.1)
Neisseria lactamica 020-06 (YP_004049219.1)
Achromobacter xylosoxidans A8 (YP_003980578.1)
Bordetella petrii DSM 12804 (YP_001632957.1)
Mesorhizobium sp. B2-1-8 (WP_140764317.1)
Mesorhizobium sp. N.Ca.IN.012.08.1 (RWM33537.1)
Acidisphaera sp. S103 (WP_158927615.1)
Uncultured Hyphomicrobiales bacterium (MBN9078826.1)
Bradyrhizobium canariense GAS369 (WP_100383287.1)
Geminicoccus sp. CPCC 101083 (WP_191060274.1)
Rhodovastum atsumiense G2-11 (WP_150040348.1)
Rhodopila globiformis DSM 161 (WP_104520311.1)
Rhodoblastus acidophilus (MBB4201663.1 )
Uncultured Acetobacteraceae bacterium HOG41 (MBX5471092.1)
Methylocella tundrae T4 WP 134491896.1
Methylocella tundrae PC (VTZ21898.1)
Methylocella tundrae PC4 (WP_174512061.1)


122

**Fig. S1.** Phylogenetic reconstruction of NosZ proteins encoded in methanotrophs and other prokaryotes. A maximum-likelihood tree was inferred with IQ-TREE using ModelFinderPlus (IQ-TREE options: -m MFP, -B 1000) and rooted at the mid-point. Methanotrophs with clade I and clade II NosZ proteins have their names written in red and blue text, respectively. Bootstrap values higher than 70% are indicated. The scale bar represents a 0.5 change in each amino acid position.

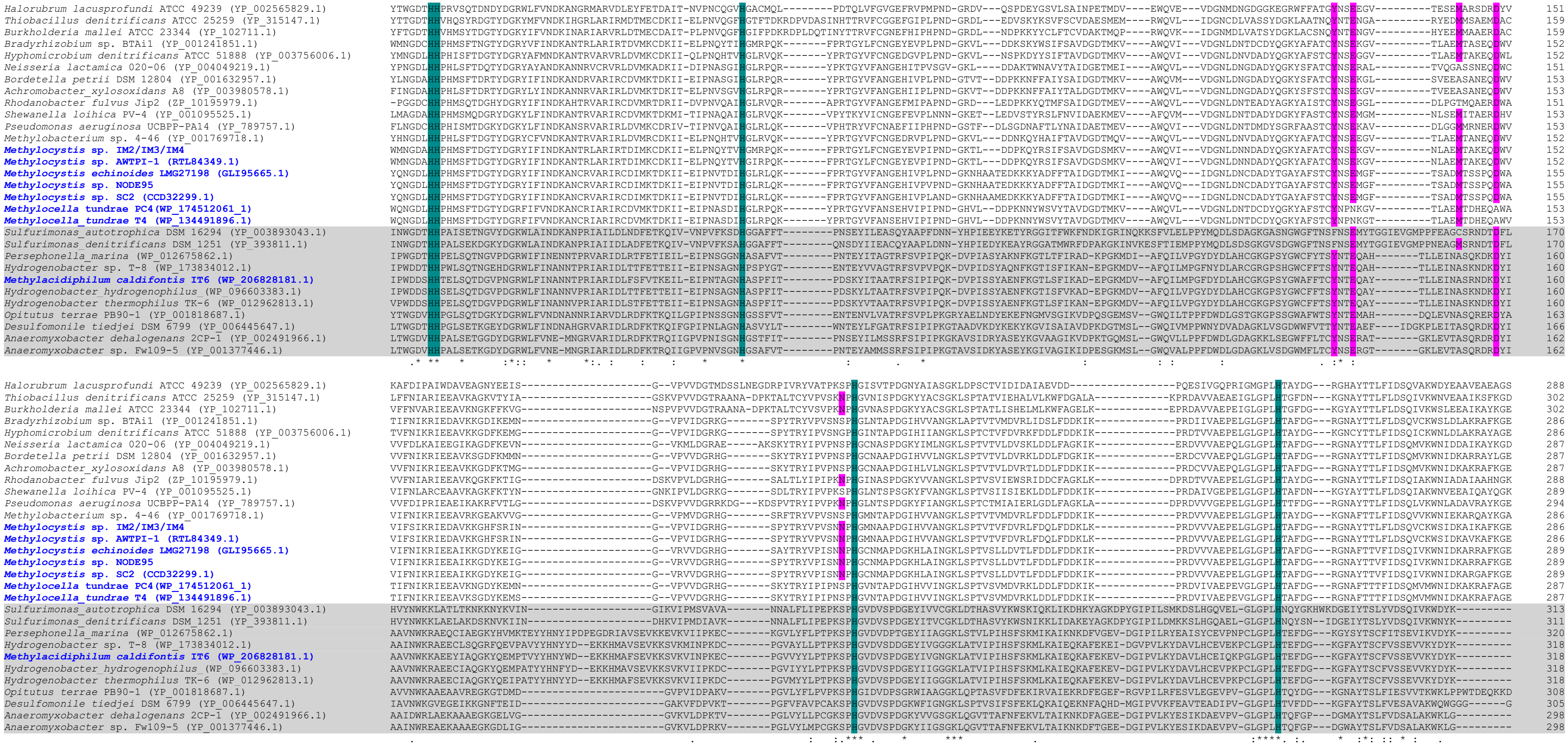

Halorubrum lacusprofundi ATCC 49239 (YP_002565829.1)
Thiobacillus denitrificans ATCC 25259 (YP_315147.1)
Burkholderia mallei ATCC 23344 (YP_102711.1)
Bradyrhizobium sp. BTAi1 (YP_001241851.1)
Hyphomicrobium denitrificans ATCC 51888 (YP_003756006.1)
Neisseria lactamica 020-06 (YP_004049219.1)
Bordetella petrii DSM 12804 (YP_001632957.1)
Achromobacter_xylosoxidans A8 (YP_003980578.1)
Rhodanobacter fulvus Jip2 (ZP_10195979.1)
Shewanella loihica PV-4 (YP_001095525.1)
Pseudomonas aeruginosa UCBPP-PA14 (YP_789757.1)
Methylobacterium sp. 4-46 (YP_001769718.1)
Methylocystis sp. IM2/IM3/IM4
Methylocystis sp. AWTPI-1 (RTL84349.1)
Methylocystis echinoides LMG27198 (GLI95665.1)
Methylocystis sp. NODE95
Methylocystis sp. SC2 (CCD32299.1)
Methylocella tundrae PC4 (WP_174512061_1)
Methylocella_tundrae T4 (WP_134491896.1)
Sulfurimonas_autotrophica DSM 16294 (YP_003893043.1)
Sulfurimonas_denitrificans DSM 1251 (YP_393811.1)
Persephonella_marina (WP_012675862.1)
Hydrogenobacter sp. T-8 (WP_173834012.1)
Methylacidiphilum caldifontis IT6 (WP_206828181.1)
Hydrogenobacter_hydrogenophilus (WP_096603383.1)
Hydrogenobacter thermophilus TK-6 (WP_012962813.1)
Opitutus terrae PB90-1 (YP_001818687.1)
Desulfomonile tiedjei DSM 6799 (YP_006445647.1)
Anaeromyxobacter dehalogenans 2CP-1 (YP_002491966.1)
Anaeromyxobacter sp. Fw109-5 (YP_001377446.1)

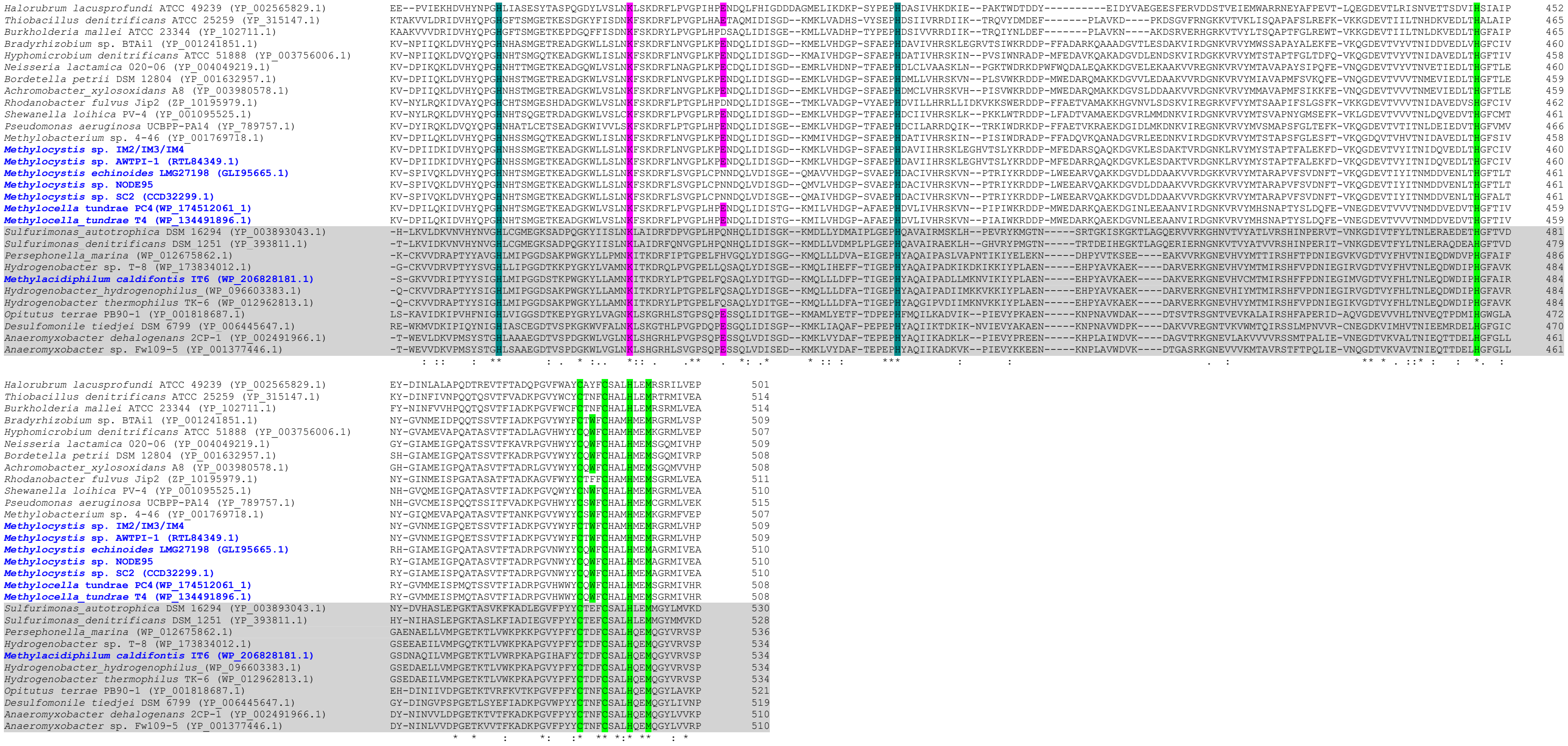

**Fig. S2**. Sequence conservation across NosZ homologues. Conserved histidine residues of the beta-propeller domain implicated in $Cu_Z$-center coordination are highlighted in teal green. Residues involved in binding the $Cu_A$-center of the C-terminal cupredoxin domain are highlighted in bright green. Calcium binding residues, involved in NosZ dimerization, are highlighted in pink. Amino acid conservation between the proteins is indicated below the alignments with the following symbols in parentheses: (*****), 100% identical amino acid residues; (**:**), conserved substitutions; (**.**), semi-conserved substitutions. NosZ proteins highlighted in grey are clade II type, while unhighlighted are clade I type. Names of NosZ-containing methanotrophs are written in blue text.

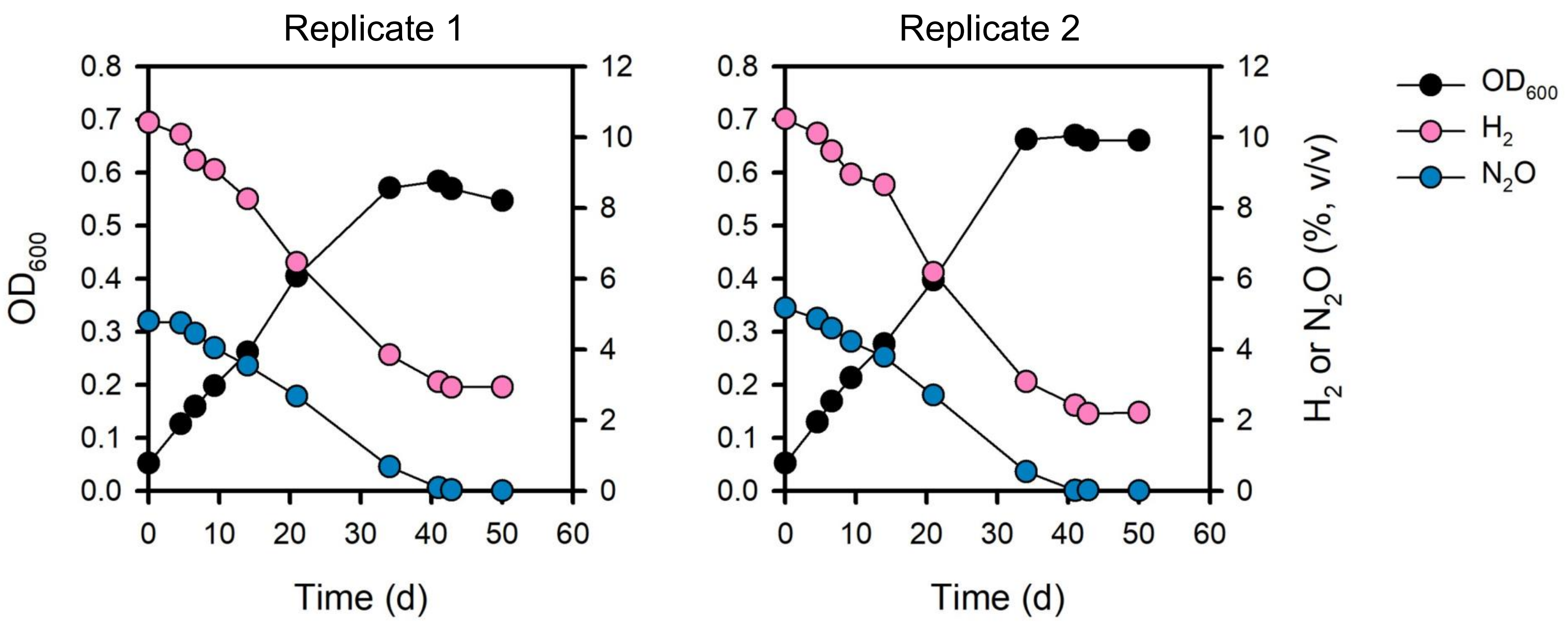


**Fig. S3.** Anaerobic growth of *Methylacidiphilum caldifontis* IT6 on hydrogen coupled with $N_2O$ reduction. *Methylacidiphilum caldifontis* IT6 cells were grown in 1-liter bottles (2 replicates) containing 60 mL of LSM medium at pH 2.0 and a headspace containing 10% (v/v) $H_2$ as an electron donor, 5% (v/v) $N_2O$ as an electron acceptor, and 5% (v/v) $CO_2$ as a carbon source. Optical density measurements at 600 nm were used to determine growth, followed by $H_2$ and $N_2O$ consumption measurements in the culture bottles' headspace.

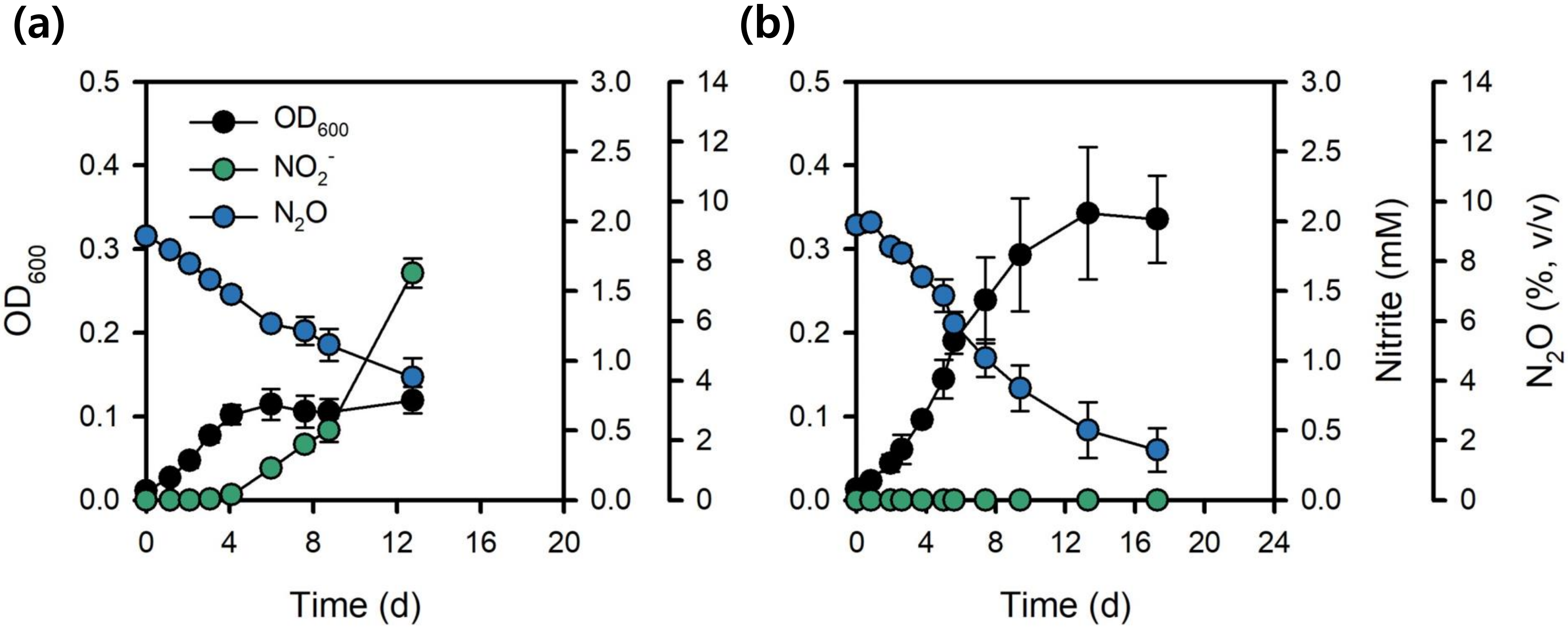


**Fig. S4.** The effect of $NO_2^-$ accumulation on *Methylocella tundrae* T4 anaerobic growth on $CH_3OH$ and $N_2O$. Strain T4 cells were grown anaerobically on $CH_3OH$ and $N_2O$ in $NO_3^-$-containing (a) and $NO_3^-$-free (b) media. $NO_2^-$ formed by the reduction of $NO_3^-$ inhibits anaerobic growth in the $NO_3^-$-containing medium. Growth inhibition was not shown in the $NO_3^-$-free medium. Data from **a** and **b** are the means of three biological replicates ± SD. The error bars are hidden when they are smaller than the width of the symbols' size.

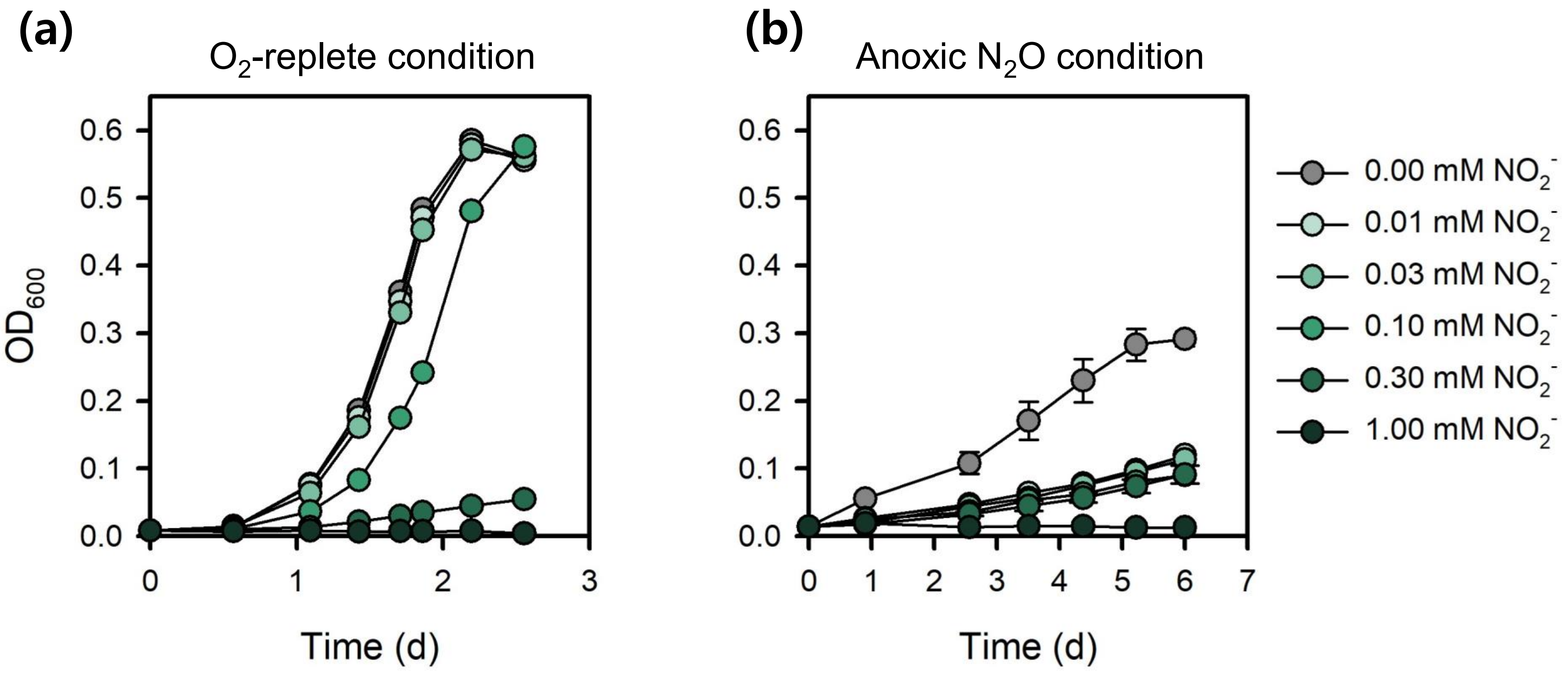


**Fig. S5**. Aerobic and anaerobic growth of *Methylocella tundrae* T4 on $CH_3OH$ in response to different $NO_2^-$ concentrations. Cultures of strain T4 grown on methanol under (**a**) $O_2$-replete and (**b**) anoxic $N_2O$-respiring growth conditions were incubated with different initial concentrations of $NO_2^-$. The presence of $NO_2^-$ was inhibitory to both conditions. Data from **a** and **b** are the means of three biological replicates. The error bars are hidden when they are smaller than the width of the symbols' size.

*M. tundrae* T4

*M.* caldifontis IT6

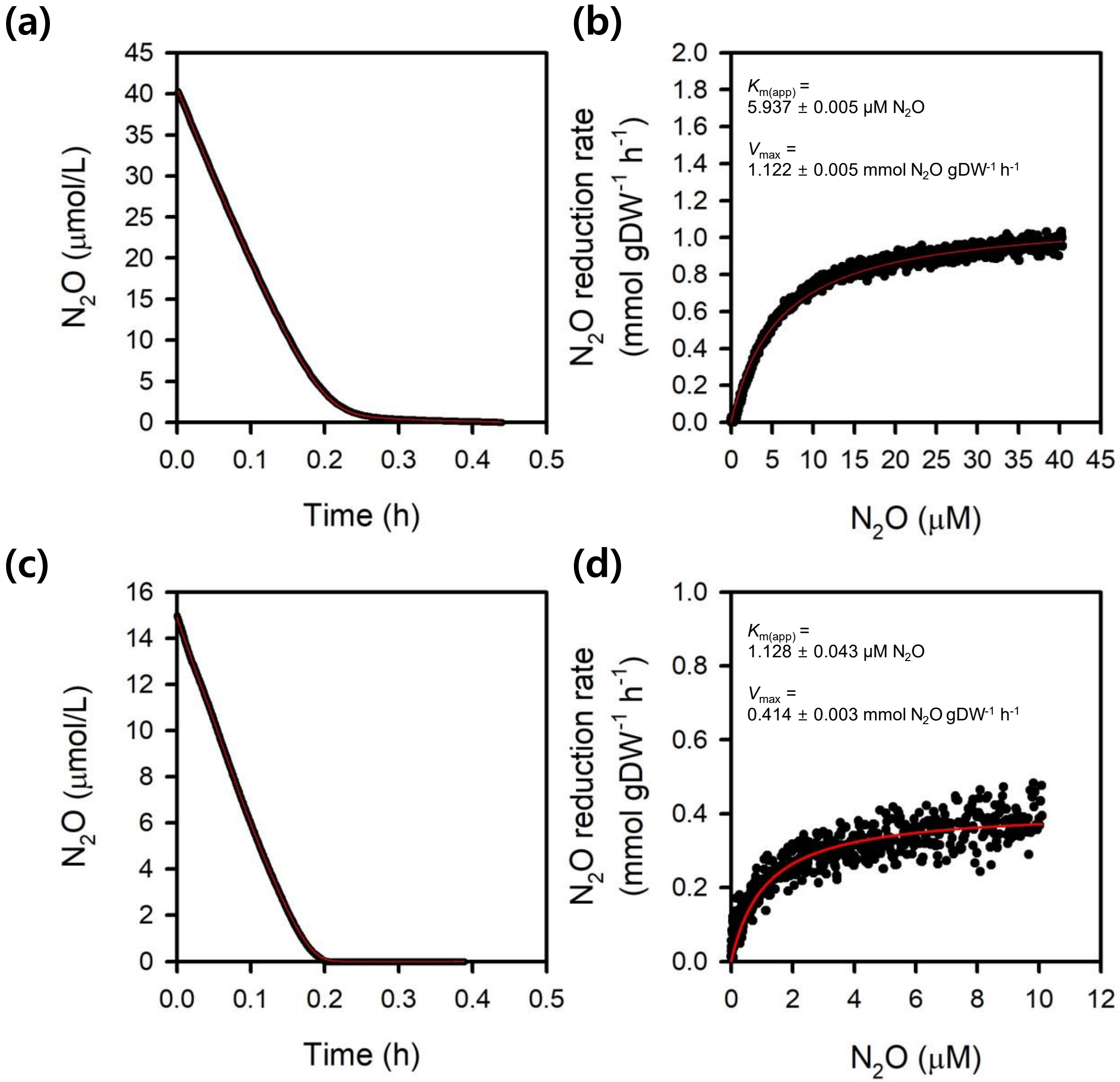

**Fig. S6.** $N_2O$ reduction kinetics of strains T4 and IT6. (**a**) $N_2O$ reduction by anoxic $CH_3OH$ + $N_2O$-grown *Methylocella tundrae* T4 cells. (**b**) Michaelis–Menten plot of $N_2O$ reduction by anoxic $CH_3OH$ + $N_2O$-grown T4 cells. (**c**) $N_2O$ reduction by anoxic $CH_3OH$ + $N_2O$-grown *Methylacidiphilum caldifontis* IT6 cells. (**d**) Michaelis–Menten plot of $N_2O$ reduction by anoxic $CH_3OH$ + $N_2O$-grown IT6 cells. The $N_2O$ reduction rate was determined from microsensor measurements of methanol-dependent $N_2O$ reduction from a single trace measurement. The Michaelis-Menten kinetic equation was fitted to the data to determine the apparent half-saturation ($K_{m(app)}$) and maximum $N_2O$ reduction rates ($V_{max}$). The red line represents the best fit to the data. The standard deviations of the non-linear regression estimates are given.

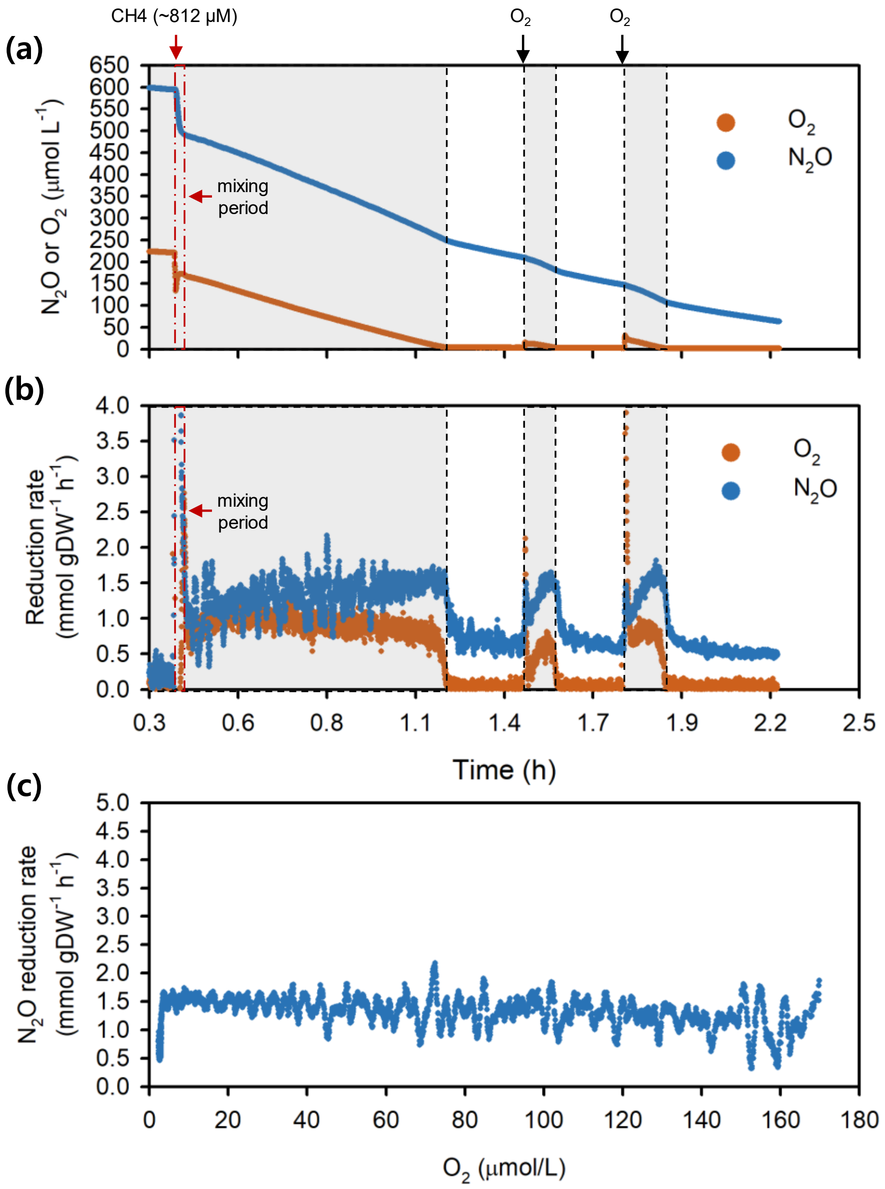


**Fig. S7.** $CH_4$-dependent $N_2O$ reduction by *Methylocella tundrae* T4 cells at high $O_2$ concentrations. (**a**) Microrespirometry experiment demonstrating $CH_4$-dependent $N_2O$ reduction at high DO concentrations by *Methylocella tundrae* T4 cells. (**b**) $N_2O$ and $O_2$ reduction rates by cells of strain T4 during $CH_4$ oxidation were calculated from the upper panel (**a**). The orange and blue dots in the upper panel (**a**) represent the concentrations of dissolved $O_2$ and $N_2O$, respectively. The orange and blue dots in the bottom panel (**b**) represent the rates of $O_2$ and $N_2O$ reduction, respectively. (**c**) $N_2O$ reduction rates at varying $O_2$ concentrations are calculated from the upper panel (**a**). Experiments were performed in a microrespiration (MR) chamber fitted with $O_2$ and $N_2O$ microsensors. The red and black arrows mark the addition of $CH_4$ (~ 812 µM) and $O_2$ into the MR chamber, respectively. The gray-shaded area represents periods where $N_2O$ and $O_2$ are reduced simultaneously.

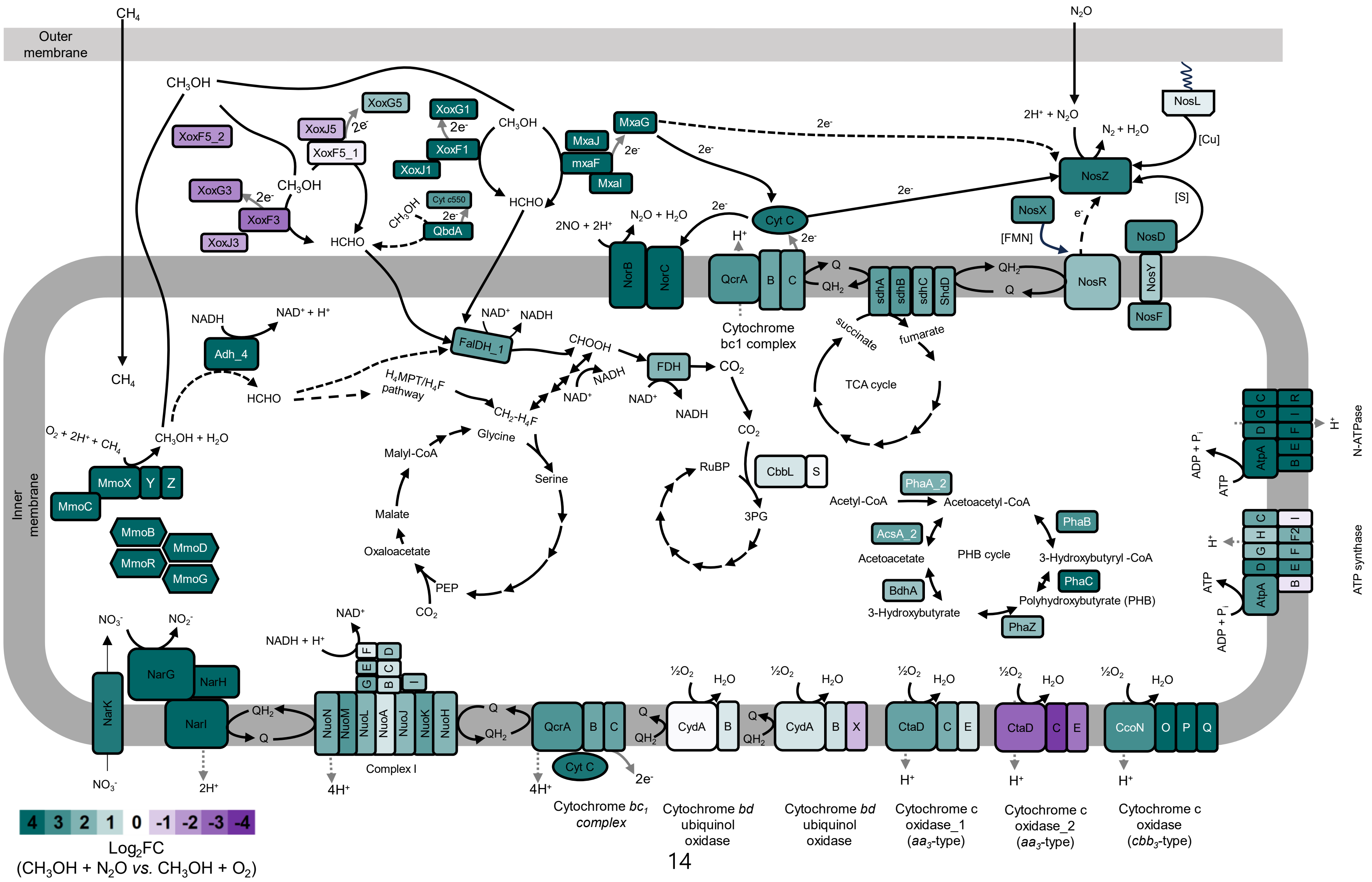

Outer membrane
Inner membrane
$CH_4$
$CH_3OH$
$N_2O$
$2H^+ + N_2O$
$N_2 + H_2O$
NosL
[Cu]
NosZ
[S]
NosX
[FMN]
NosD
NosR
NosY
NosF
MxaG
MxaJ
mxaF
MxaI
XoxG5
XoxJ5
XoxF5_1
XoxF5_2
XoxG1
XoxF1
XoxJ1
XoxG3
XoxF3
XoxJ3
Cyt c550
QbdA
HCHO
Cyt C
NorB
NorC
$2NO + 2H^+$
$N_2O + H_2O$
QcrA
Cytochrome bc1 complex
sdhA
sdhB
sdhC
ShdD
succinate
fumarate
TCA cycle
NADH
$NAD^+ + H^+$
Adh_4
$H_4MPT/H_4F$ pathway
FalDH_1
CHOOH
FDH
$CO_2$
$CH_2$-$H_4F$
Glycine
Serine
Malyl-CoA
Malate
Oxaloacetate
PEP
RuBP
3PG
CbbL
S
Acetyl-CoA
Acetoacetyl-CoA
PhaA_2
PhaB
AcsA_2
Acetoacetate
PHB cycle
3-Hydroxybutyryl -CoA
PhaC
BdhA
3-Hydroxybutyrate
Polyhydroxybutyrate (PHB)
PhaZ
$O_2 + 2H^+ + CH_4$
$CH_3OH + H_2O$
MmoX
Y
Z
MmoC
MmoB
MmoD
MmoR
MmoG
AtpA
N-ATPase
ATP synthase
$ADP + P_i$
ATP
$NO_3^-$
$NO_2^-$
NarK
NarG
NarH
NarI
$QH_2$
Q
2H+
NADH + H+
NAD+
NuoN
NuoM
NuoL
NuoA
NuoJ
NuoK
NuoH
Complex I
4H+
Cytochrome $bc_1$ complex
CydA
Cytochrome bd ubiquinol oxidase
½$O_2$
$H_2O$
CtaD
Cytochrome c oxidase_1 ($aa_3$-type)
Cytochrome c oxidase_2 ($aa_3$-type)
CcoN
Cytochrome c oxidase ($cbb_3$-type)
4 3 2 1 0 -1 -2 -3 -4
$Log_2FC$
($CH_3OH + N_2O$ vs. $CH_3OH + O_2$)

**Fig. S8.** Metabolic reconstruction and transcriptional response of methane-oxidizing *Methylocella tundrae* strain T4 cells to $O_2$-replete $CH_4 + O_2$- and suboxic $CH_4 + O_2 + N_2O$-growth conditions. The genes used to reconstruct the metabolic pathway are listed in Table S5. The gene products are shaded according to the relative fold change ($Log_2FC$) in gene expression between methane-oxidizing cells grown in suboxic $CH_4 + O_2 + N_2O$ and $O_2$-replete $CH_4 + O_2$ conditions. Genes up-regulated in suboxic $CH_4 + O_2 + N_2O$-grown cells are shown in teal green, while those up-regulated in $O_2$-replete $CH_4 + O_2$-grown cells are shown in purple. Note that proteins are not drawn to scale. **Methane oxidation:** Methane is oxidized to methanol by the cytoplasmic (soluble) methane monooxygenase, sMMO (T4_01946–54). **Methanol oxidation**: Methanol is oxidized to formaldehyde in the periplasmic space by the PQQ-dependent methanol dehydrogenase (Xox- and Mxa-type), T4_03519-21, T4_00353-55, T4_01862-76. Methanol oxidation may also be mediated by the type II quinohemoprotein alcohol dehydrogenase (T4_02097-98). The $NAD(P)^+$-dependent alcohol dehydrogenase (T4_03199) may also be involved in methanol oxidation to formaldehyde in the cytoplasmic space during anaerobic growth on methanol. Formaldehyde oxidation to formate then proceeds via the tetrahydromethanopterin ($H_4MPT$) pathway, and C1 incorporation into the serine cycle is mediated by the tetrahydrofolate ($H_4F$) carbon assimilation pathway. The Calvin-Benson-Bassham pathway is also a possible route for $CO_2$ fixation. **Nitrous oxide reduction**: $N_2O$ is reduced to $N_2$ through the activity of $N_2OR$ in the periplasmic space. Electron transfer to NosZ occurs via cytochrome c from the cytochrome bc1 (Qcr) complex. Electron transfer to the NosZ may also involve direct interaction with methanol dehydrogenase c-type cytochrome (XoxG, MxaG). The NosR protein may be involved in the transfer of electrons to NosZ. The $N_2O$ reduction pathway is adopted from Hein and Simon (7) and Torres *et al*. (8).

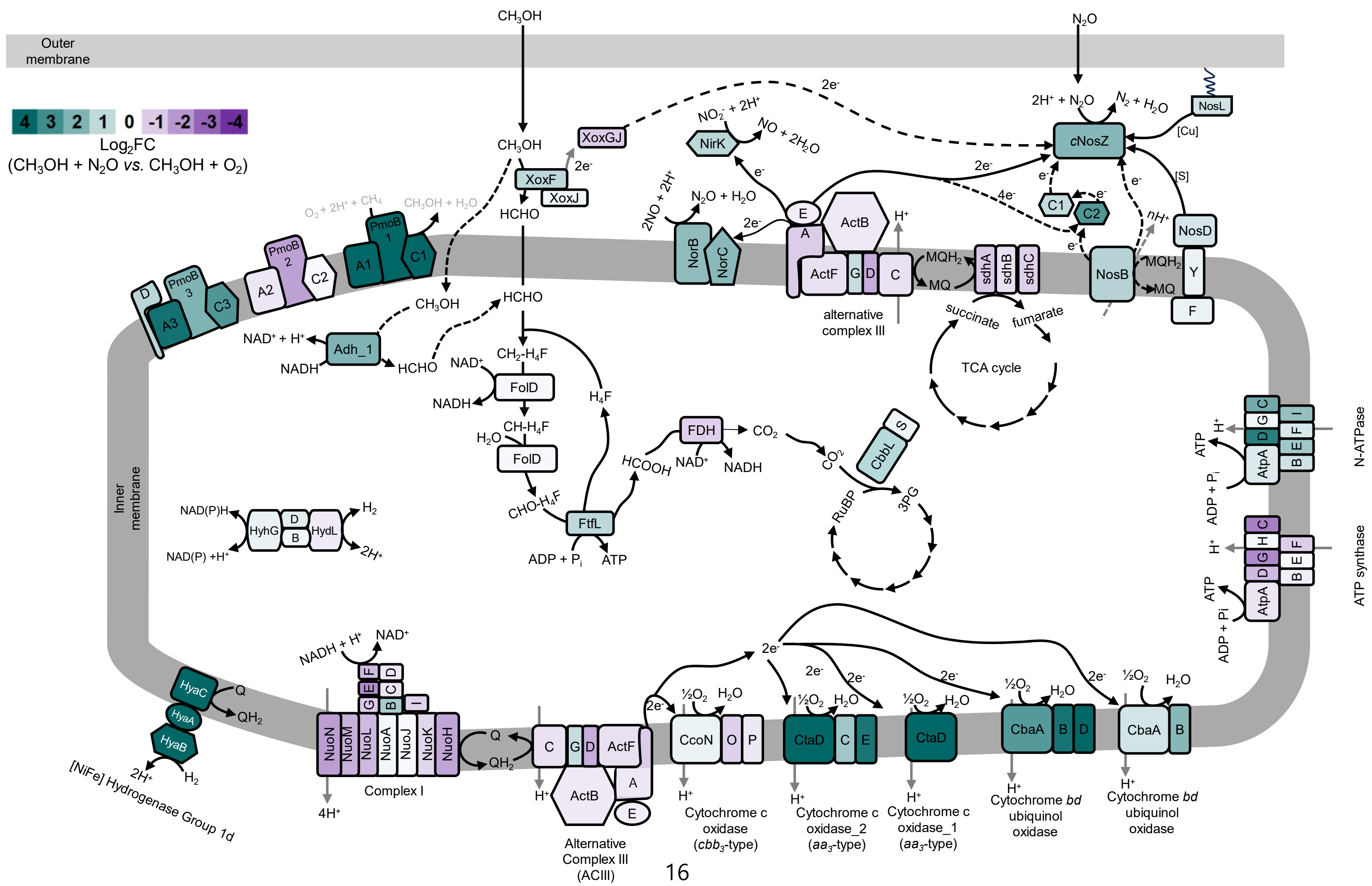

Outer membrane
Inner membrane
4 3 2 1 0 -1 -2 -3 -4
Log2FC
(CH3OH + N2O vs. CH3OH + O2)
CH3OH
XoxGJ
XoxF
XoxJ
2e-
HCHO
N2O
2H+ + N2O
N2 + H2O
cNosZ
NosL
[Cu]
[S]
NosD
NosB
Y
F
C1
C2
nH+
MQH2
MQ
NO2- + 2H+
NirK
NO + 2H2O
N2O + H2O
2NO + 2H+
NorB
NorC
ActB
ActF
alternative complex III
sdhA
sdhB
sdhC
succinate
fumarate
TCA cycle
O2 + 2H+ + CH4
CH3OH + H2O
PmoB 1
PmoB 2
PmoB 3
A1
A2
A3
C1
C2
C3
D
Adh_1
NAD+ + H+
NADH
CH2-H4F
FolD
NAD+
NADH
CH-H4F
H2O
CHO-H4F
FtfL
ADP + Pi
ATP
H4F
FDH
CO2
HCOOH
CbbL
S
RuBP
3PG
NAD(P)H
NAD(P) +H+
HyhG
HydL
H2
2H+
N-ATPase
ATP synthase
AtpA
HyaC
HyaA
HyaB
Q
QH2
[NiFe] Hydrogenase Group 1d
NADH + H+
NAD+
NuoN
NuoM
NuoL
NuoA
NuoJ
NuoK
NuoH
Complex I
4H+
Alternative Complex III (ACIII)
½O2
H2O
CcoN
Cytochrome c oxidase (cbb3-type)
CtaD
Cytochrome c oxidase_2 (aa3-type)
Cytochrome c oxidase_1 (aa3-type)
CbaA
Cytochrome bd ubiquinol oxidase

**Fig. S9.** Metabolic reconstruction and transcriptional response of methanol-oxidizing *Methylacidiphilum caldifontis* IT6 cells in $O_2$-replete $CH_3OH$ + $O_2$- and anoxic $CH_3OH$ + $N_2O$-growth conditions. The genes used to reconstruct the metabolic pathway are listed in Table S6. The gene products are shaded according to the relative fold change ($Log_2FC$) in gene expression between methane-oxidizing cells grown in anoxic $CH_3OH$ + $N_2O$ and $O_2$-replete $CH_3OH$ + $O_2$ conditions. Genes up-regulated in anoxic $CH_3OH$ + $N_2O$-grown cells are shown in teal green, while those up-regulated in $O_2$-replete $CH_3OH$ + $O_2$-grown cells are shown in purple. Note that proteins are not drawn to scale. **Methanol oxidation**: Methanol is oxidized to formaldehyde in the periplasmic space by the PQQ-dependent methanol dehydrogenase (Xox-type), IT6_00336-38. The $NAD(P)^+$-dependent alcohol dehydrogenase (IT6_01501 and IT6_01931) may also be involved in methanol oxidation to formaldehyde in the cytoplasmic space during anaerobic growth on methanol. Meth anol oxidation produces formaldehyde, which can either spontaneously or enzymatically bind to tetrahydrofolate ($H_4F$) to produce methylene-tetrahydrofolate ($CH_2$-$H_4F$). FolD converts methylenetrahydrofolate ($CH_2$-$H_4F$) to methenyl-tetrahydrofolate (CH-$H_4F$), which is then converted to formyl-tetrahydrofolate (CHO-$H_4F$) by the same enzyme. ATP is generated in the process of converting this product to $H_4F$ and formate. Formate is oxidized to $CO_2$ through the activity of formate dehydrogenase. The pathway for methanol oxidation to $CO_2$ is adopted from Schmitz *et al*. (9). C1 incorporation into biomass is through the Calvin-Benson-Bassham pathway. **Nitrous oxide reduction**: Strain IT6 produces a cytochrome *c* $N_2OR$ (*c*NosZ) with an additional C-terminal monohaem cytochrome *c* domain thought to function as an electron entry point to the active site copper. The alternative complex III (ACIII) oxidizes $MQH_2$ to MQ and transfers electrons to NosZ. The methanol dehydrogenase *c*-type cytochrome (XoxGJ) may also be involved in electron transfer to the NosZ. Electron transfer from menaquinol to NosZ is predicted to involve NosB, NosC1, and NosC2. The $N_2O$ reduction pathway and the Nos protein localization are adapted from Simon *et al*. (10), Hein and Simon (7), and Torres *et al*. (8).

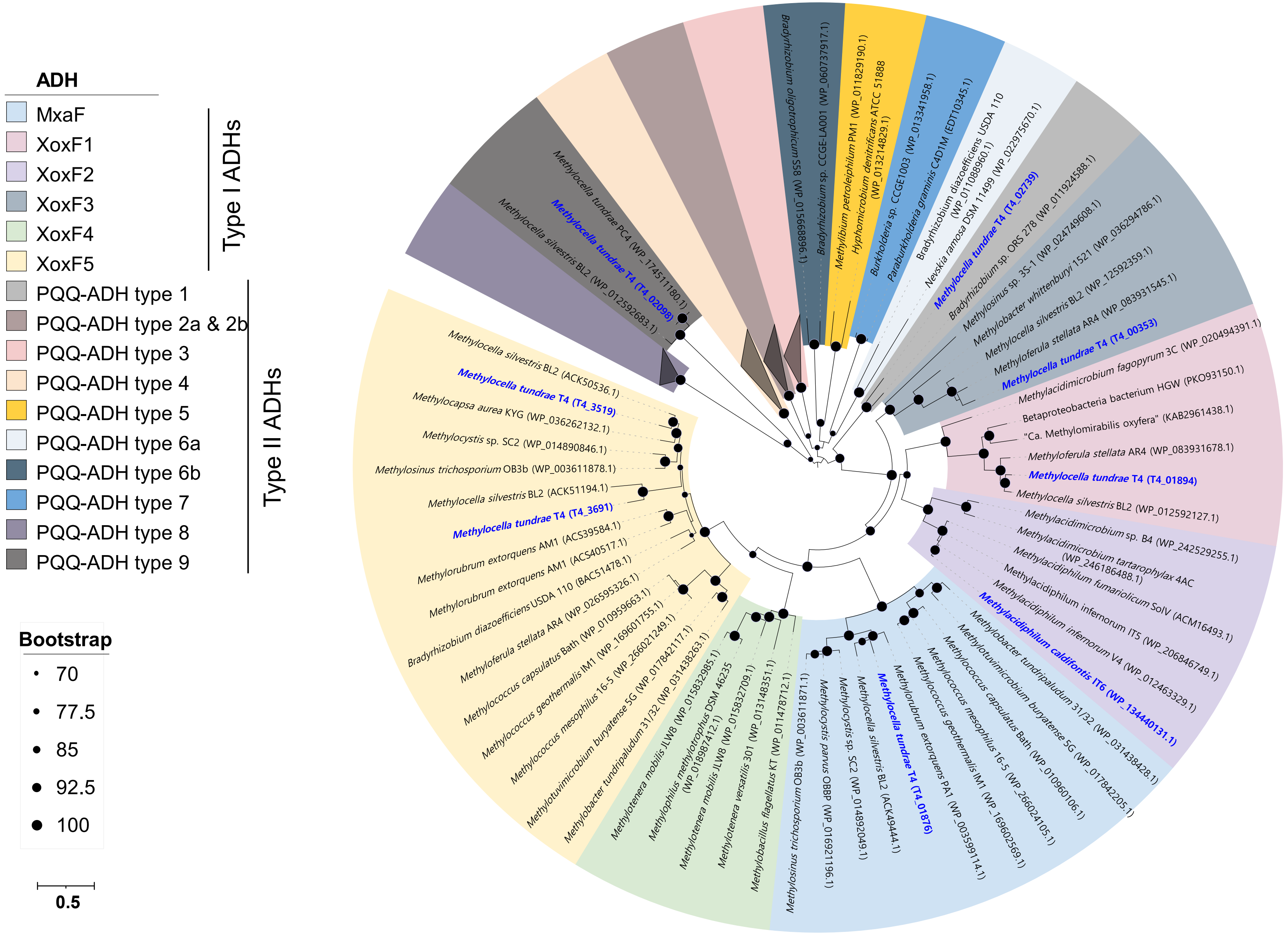

ADH
MxaF
XoxF1
XoxF2
XoxF3
XoxF4
XoxF5
PQQ-ADH type 1
PQQ-ADH type 2a & 2b
PQQ-ADH type 3
PQQ-ADH type 4
PQQ-ADH type 5
PQQ-ADH type 6a
PQQ-ADH type 6b
PQQ-ADH type 7
PQQ-ADH type 8
PQQ-ADH type 9
Type I ADHs
Type II ADHs
Bootstrap
70
77.5
85
92.5
100
0.5
Methylocella silvestris BL2 (WP_012592683.1)
Methylocella tundrae T4 (T4_02098)
Methylocella tundrae PC4 (WP_174511180.1)
Bradyrhizobium oligotrophicum S58 (WP_015668896.1)
Bradyrhizobium sp. CCGE-LA001 (WP_060737917.1)
Methylibium petroleiphilum PM1 (WP_011829190.1)
Hyphomicrobium denitrificans ATCC 51888 (WP_013214829.1)
Burkholderia sp. CCGE1003 (WP_013341958.1)
Paraburkholderia graminis C4D1M (EDT10345.1)
Bradyrhizobium diazoefficiens USDA 110 (WP_011088960.1)
Nevskia ramosa DSM 11499 (WP_022975670.1)
Methylocella tundrae T4 (T4_02739)
Bradyrhizobium sp. ORS 278 (WP_011924588.1)
Methylosinus sp. 3S-1 (WP_024749608.1)
Methylobacter whittenburyi 1521 (WP_036294786.1)
Methylocella silvestris BL2 (WP_12592359.1)
Methyloferula stellata AR4 (WP_083931545.1)
Methylocella tundrae T4 (T4_00353)
Methylacidimicrobium fagopyrum 3C (WP_020494391.1)
Betaproteobacteria bacterium HGW (PKO93150.1)
"Ca. Methylomirabilis oxyfera" (KAB2961438.1)
Methyloferula stellata AR4 (WP_083931678.1)
Methylocella tundrae T4 (T4_01894)
Methylocella silvestris BL2 (WP_012592127.1)
Methylacidimicrobium sp. B4 (WP_242529255.1)
Methylacidimicrobium tartarophylax 4AC (WP_246186488.1)
Methylacidiphilum fumariolicum SolV (ACM16493.1)
Methylacidiphilum infernorum ITS (WP_206846749.1)
Methylacidiphilum infernorum V4 (WP_012463329.1)
Methylacidiphilum caldifontis IT6 (WP_134440131.1)
Methylotuvimicrobium buryatense 5G (WP_031438428.1)
Methylobacter tundripaludum 31/32 (WP_017842205.1)
Methylococcus capsulatus Bath (WP_010960106.1)
Methylococcus mesophilus 16-5 (WP_266024105.1)
Methylococcus geothermalis IM1 (WP_169602569.1)
Methylorubrum extorquens PA1 (WP_003599114.1)
Methylocella tundrae T4 (T4_01876)
Methylocella silvestris BL2 (ACK49444.1)
Methylocystis sp. SC2 (WP_014892049.1)
Methylocystis parvus OBBP (WP_016921196.1)
Methylosinus trichosporium OB3b (WP_003611871.1)
Methylobacillus flagellatus KT (WP_011478712.1)
Methylotenera versatilis 301 (WP_013148351.1)
Methylotenera mobilis JLW8 (WP_015832709.1)
Methylophilus methylotrophus DSM 46235 (WP_018987412.1)
Methylotenera mobilis JLW8 (WP_015832985.1)
Methylobacter tundripaludum 31/32 (WP_031438263.1)
Methylotuvimicrobium buryatense 5G (WP_017842117.1)
Methylococcus mesophilus 16-5 (WP_266021249.1)
Methylococcus geothermalis IM1 (WP_169601755.1)
Methylococcus capsulatus Bath (WP_010959663.1)
Methyloferula stellata AR4 (WP_026595326.1)
Bradyrhizobium diazoefficiens USDA 110 (BAC51478.1)
Methylorubrum extorquens AM1 (ACS40517.1)
Methylorubrum extorquens AM1 (ACS39584.1)
Methylocella tundrae T4 (T4_3691)
Methylocella silvestris BL2 (ACK51194.1)
Methylosinus trichosporium OB3b (WP_003611878.1)
Methylocystis sp. SC2 (WP_014890846.1)
Methylocapsa aurea KYG (WP_036262132.1)
Methylocella tundrae T4 (T4_3519)
Methylocella silvestris BL2 (ACK50536.1)

**Fig. S10**. Phylogenetic analysis of PQQ-dependent alcohol dehydrogenases (type I quinoproteins and type II quinohemoprotein). Different subclasses or clades of the type I quinoproteins and type II quinohemoprotein are shown. A maximum-likelihood tree was inferred with IQ-TREE (IQ-TREE options: -B 1000 -m LG+F+R5 -T AUTO) and rooted at the mid-point. Bootstrap values higher than 70% are indicated. PQQ-dependent ADH of strains T4 and IT6 are labeled blue.

**Table S1 (separate file).**

Denitrification genes distribution in methanotrophs. Isolates and metagenome-assembled genomes of methanotrophs are included. Genes encoding the soluble and particulate methane monooxygenases were included. The presence of each gene in target genomes is denoted by the green color bar and the number of copies is included within the bar. Published genomes and MAGs were compiled from the NCBI database.

**Table S2 (separate file).**

The *nos* gene cluster (NGC) of methanotrophs. The highest percent similarity with the translated amino acid sequence of the genes of NGC and entries in the NCBI database is shown.

**Table S3 (separate file).**

Genomic island-related genes found in the genome of *Methylacidiphilum caldifontis* IT6. The genomic islands were identified with IslandViewer 4.

**Table S4 (separate file).**

$N_2O$-dependent anaerobic growth of $N_2OR$-containing and $N_2OR$-lacking strains of genera *Methylocella* and *Methylacidiphilum* under different electron donors. Growth is indicated as: + if the increase in $OD_{600}$ > 0.02; − if the increase in $OD_{600}$ < 0.005. $OD_{600}$ < 0.005; ND, not determined. All cultures were cultivated in LSM media at pH 2 (*Methylacidiphilum* spp.) and pH 5.5 (*Methylocella* spp.).

**Table S5 (separate file).**

Central metabolic genes in *Methylocella tundrae* T4 and their differential expression under methanol- ($CH_3OH$ + $N_2O$ versus $CH_3OH$ + $O_2$) and methane- ($CH_4$ + $O_2$ + $N_2O$ versus $CH_4$ + $O_2$) oxidizing growth conditions. Differences in expression were considered upregulated if the $Log_2FC$ was higher than [0.85] or downregulated if lower than [-1.0] with an adjusted *p*-value ≤ 0.05. Data from four biological replicates. For easy comparisons between samples, TPM (Transcripts Per Kilobase Million) values were calculated.

**Table S6 (separate file).**

Central metabolic genes in *Methylacidiphilum caldifontis* IT6 and their differential expression under anoxic $CH_3OH$ + $N_2O$ *vs.* $O_2$-replete $CH_3OH$ + $O_2$-growth conditions. Differences in expression were considered upregulated if the $Log_2FC$ was higher than [0.85] or downregulated if lower than [-1.0] with an adjusted p-value ≤ 0.05. Data from four or five biological replicates. For easy comparisons between samples, TPM (Transcripts Per Kilobase Million) values were calculated.
